# Phage predation, disease severity and pathogen genetic diversity in cholera patients

**DOI:** 10.1101/2023.06.14.544933

**Authors:** Naïma Madi, Emilee T. Cato, Md. Abu Sayeed, Ashton Creasy-Marrazzo, Aline Cuénod, Kamrul Islam, Md. Imam UL. Khabir, Md. Taufiqur R. Bhuiyan, Yasmin A. Begum, Emma Freeman, Anirudh Vustepalli, Lindsey Brinkley, Manasi Kamat, Laura S. Bailey, Kari B. Basso, Firdausi Qadri, Ashraful I. Khan, B. Jesse Shapiro, Eric J. Nelson

## Abstract

Despite an increasingly detailed picture of the molecular mechanisms of phage-bacterial interactions, we lack an understanding of how these interactions evolve and impact disease within patients. Here we report a year-long, nation-wide study of diarrheal disease patients in Bangladesh. Among cholera patients, we quantified *Vibrio cholerae* (prey) and its virulent phages (predators) using metagenomics and quantitative PCR, while accounting for antibiotic exposure using quantitative mass spectrometry. Virulent phage (ICP1) and antibiotics suppressed *V. cholerae* to varying degrees and were inversely associated with severe dehydration depending on resistance mechanisms. In the absence of anti-phage defenses, predation was ‘effective,’ with a high predator:prey ratio that correlated with increased genetic diversity among the prey. In the presence of anti-phage defenses, predation was ‘ineffective,’ with a lower predator:prey ratio that correlated with increased genetic diversity among the predators. Phage-bacteria coevolution within patients should therefore be considered in the deployment of phage-based therapies and diagnostics.

**One Sentence Summary:** A survey of cholera patients in Bangladesh identifies phage predation as a biomarker of disease severity and driver of coevolution within patients.

## Introduction

Cholera is caused by the Gram negative bacterium *V. cholerae* (*Vc*) and can progress to life-threatening hypovolemic shock in less than 12 hours (*1*). Cholera remains a major public health problem because of inadequate sanitation and restricted access to safe drinking water. Global estimates of the cholera burden are 1.3-4.0 million cases and 21,000-143,000 deaths annually (*2*). In 2023, there were over 30 countries with active outbreaks necessitating the WHO to escalate the response to its highest level (*3*). Rehydration is the primary life-saving intervention for cholera patients. With adequate rehydration, mortality rates fall from over 20% to less than 1%. Antibiotics reduce stool volume and duration of diarrhea but are generally reserved for patients with more severe disease to reduce selection for antibiotic resistance (*4–8*). Nevertheless, antibiotic-resistant *Vc* have emerged globally (*6, 9, 10*). Mechanisms of resistance are diverse and reside in the core genome, plasmids of the incompatibility type C (*11, 12*) and on mobile genetic elements, including a ∼100kb integrative conjugative element (ICE), which can harbor resistance to sulfamethoxazole and trimethoprim, ciprofloxacin (*qnr_vc_*), trimethoprim (*dfra31*) and streptomycin (*aph*(*6*)) (*13–16*). Recent work has also shown that the ICE can encode diverse phage resistance mechanisms, with distinct hotspots of gene content variation encoding different resistance genes (*17*).

With rising levels of antibiotic resistance, virulent bacteriophages (phages) are a promising alternative or complementary therapy to antibiotics. Phages and bacteria coevolve, with each partner selecting for adaptations in the other that generates genetic diversity for both predator and prey (*18, 19*). Coevolution likely explains the extensive arsenal of resistance and counter-resistance mechanisms among *Vc* and its phages (*17, 19–24*). Yet it remains unclear how these interactions impact disease severity during natural infection, with and without antibiotic exposure. Virulent phages specifically targeting *Vc* include ICP1 (*Myoviridae*), ICP2 (*Podoviridae*), and ICP3 (*Podoviridae*) (*21, 25, 26*). These common phages are found in symptomatic and asymptomatic cholera patients during acute infection or the convalescent period.

The first clinical trials of phage therapy occurred during the Cholera Phage Inquiry from 1927 to 1936 (*27, 28*). In a *proto* randomized controlled trial, the Inquiry found the odds of mortality were reduced by 58% among those given phage therapy, with an absolute reduction in mortality of approximately 10% (*29*). Despite these early findings, there is a lack of evidence in the modern era linking phage predation with disease severity during natural infection in humans. However, indirect studies support a link. Environmentally, virulent phages in aquatic environments have been negatively correlated with cholera cases in Bangladesh over time, suggesting a role for phages in influencing epidemic dynamics (*30*). Clinically, a higher percentage of cholera patients shed virulent phages towards the end of an outbreak period (*31*), suggesting that outbreaks may collapse because of phage predation (*8*). Theoretically, models predict that phage predation can dampen cholera outbreaks (*32*). Experimentally, animal studies found phage predation was inversely associated with colonization and severity (*25, 33–35*). The key unanswered question is if and how virulent phages, antibiotics, and bacterial evolution interact to impact infection in a meaningful manner for clinicians, public health officials, and most importantly patients.

To address this question, we conducted a national prospective longitudinal study in the cholera endemic setting of Bangladesh. Stool samples were collected at hospital admission from diarrheal disease patients and screened for *Vc*, antibiotics, and cholera phages, focusing on the obligately lytic and commonly prevalent phage ICP1. Patients in this setting routinely self-medicate with antibiotics before arriving at the hospital, hence the need to measure antibiotics in stool. We hypothesized that: (i) virulent phages and antibiotics would suppress *Vc* and be inversely associated with disease severity, (ii) suppression by phage would be lifted for *Vc* encoding anti-phage genes on ICEs, (iii) phages would select for other resistance mutations in the absence of ICE-encoded resistance, and (iv) phages would be under selection to escape suppression by ICEs. We provide broad support for these hypotheses, paving the way for mechanistic experimental studies, the development of the phage:pathogen ratio as a biomarker of disease severity, and further dissection of the longer-term consequences of phage predation on pathogen evolution.

## Results

### Study overview

A total of 2574 stool samples were collected from enrolled participants admitted for diarrheal disease at seven hospitals across Bangladesh from March to December 2018; collection continued until April 2019 at one site (icddr,b). Three groups of cholera samples were analyzed: (i) *Vc* culture-positive (282/2574; 10.9%), (ii) *Vc* culture-negative but phage (ICP1,2,3) PCR-positive (127/2292; 5.5%; 80 included; 47 excluded for DNA < 1ng/µl), and (iii) a random 10% of *Vc* culture-negative and phage PCR-negative samples that were *Vc* PCR-positive (14.8%; 37/250; 27 included; 10 excluded for DNA < 1ng/µl; see **Table S1** for PCR primers). Stool metagenomes were sequenced from 88.4% of samples (344/389, with the remainder failing library preparation) from these three groups, 35% of which were from the icddr,b site. Based on metagenomic read mapping to a taxonomic database, detection rates for *Vc,* ICP1, ICP2, and ICP3 were 55%, 18%, 1% and 8%, respectively. These detection rates were supported by an analysis of *Vc* phages identified in metagenomic assemblies. As expected, the prophage encoding the cholera toxin (CTXphi) was identified in most assemblies, with ICP1 being the most prevalent obligately virulent phage. While some additional putative phages were detected, none were prevalent enough to merit further analysis (**Fig. S1**). For both *Vc* and ICP1, relative abundances in metagenomes correlated with absolute quantification by qPCR (**Fig. S2**). Five antibiotics (**Table S2**) were prioritized for detection in stool using liquid chromatography-mass spectrometry (LCMS); of these, azithromycin, ciprofloxacin, and doxycycline were quantified.

### Metagenomic correlates of disease severity and succession

At hospital admission, we expected patients to present at different stages of disease, with an ecological succession of *Vc* followed by the facultative anaerobe *Escherichia coli,* then by a flora of mostly anaerobic bacteria (*36*). We hypothesized that these stages of succession would be associated with changes in disease severity. As expected, we identified *Vc* as an indicator species of severe dehydration. *Vc* was relatively more abundant in patients with severe dehydration, while two *Bifidobacterium* species, *E. coli*, and ICP1 were indicators of mild dehydration (**Fig. 1A, Table S3**). ICP3 was an indicator of severe dehydration despite being less frequently detected in our study (28 samples with >0.1% ICP3 reads, compared to 61 samples with >0.1% ICP1). This shows that different phages can have contrasting disease associations.

**Fig. 1.**
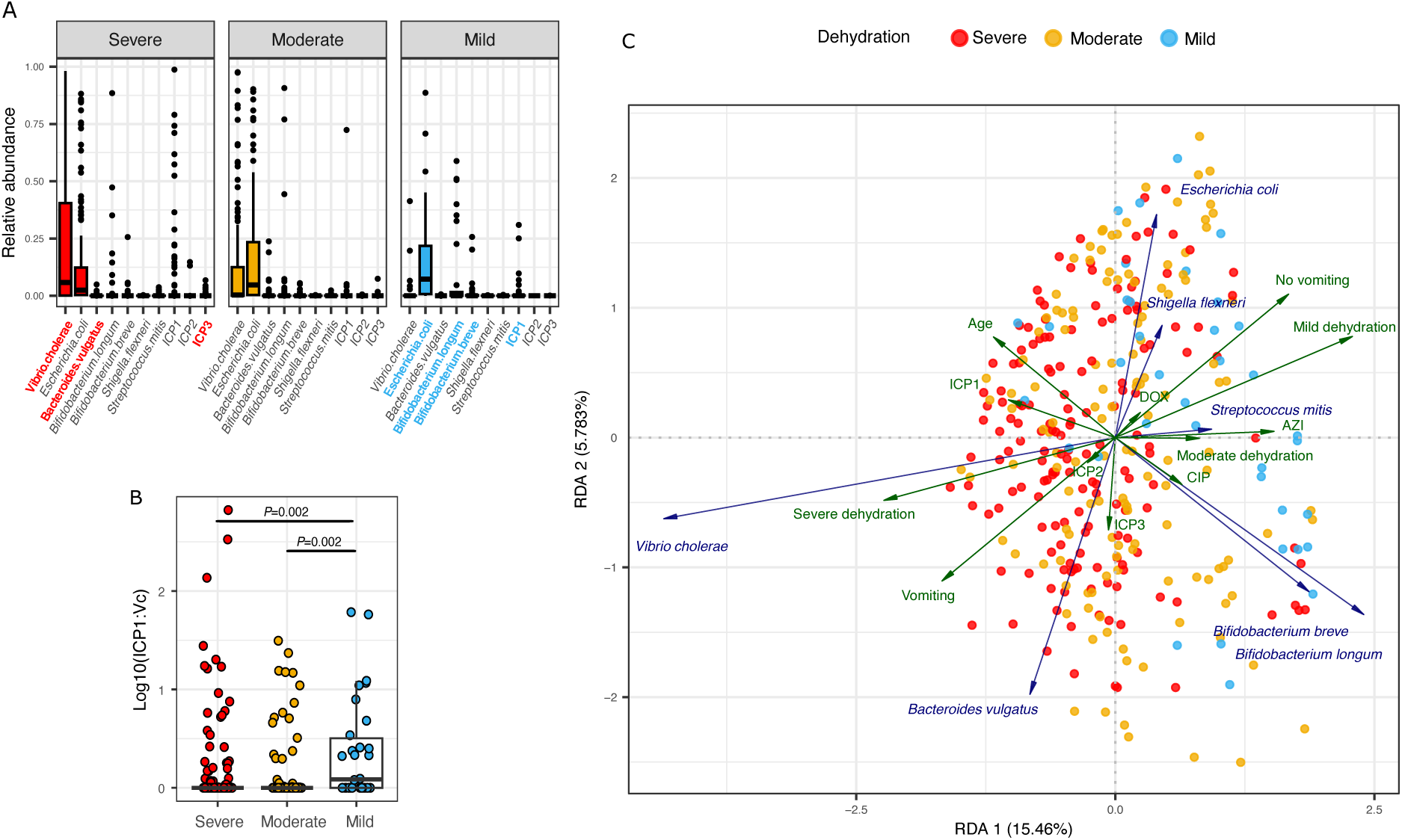
Dehydration severity is inversely associated with higher ICP1:*V. cholerae* ratios in stool metagenomes. (A) Relative abundances of phages and the seven most dominant bacterial species identified with PCA (Fig. S5) in patients with severe, moderate, or mild dehydration; these conventions equate to the World Health Organization (WHO) conventions of ‘Severe’, ‘Some’ and ‘No’ dehydration, respectively. Significant indicator species for severe or mild dehydration are shown in red or blue bold text, respectively (*P*<0.05 in a permutation test with 9999 iterations as implemented in the indicator species function in R). See Table S3 for indicator species results applied to all 37 species selected in the PCA dimensionality reduction (Fig. S5; Methods). (B) The ICP1:*Vc* ratio from metagenomics is higher in patients with mild dehydration. *P*-values are from a Kruskal-Wallis test with Dunn’s post-hoc test, adjusted for multiple tests using the Benjamini-Hochberg (BH) method. Only significant *P*-values (<0.05) are shown. Only 323 out of 344 samples were included (*Vc*>0% of metagenomic reads), with 165 from severe, 128 from moderate, and 30 from mild cases. A pseudocount of one was added to the ratio before log transformation. For supporting analyses using qPCR data, see Fig. S4. In (A) and (B) the solid horizontal line is the median and the boxed area is the interquartile range. (C) Redundancy analysis (RDA) showing relationships among the seven most dominant bacterial species identified with PCA (Fig. S5) and explanatory variables: phages (ICP1, ICP2, ICP3), patient metadata: age in years, vomiting state (yes or no), dehydration status (severe, moderate or mild), the location where the sample was collected, and date of sampling; and antibiotic concentration (µg/ml) from quantitative mass spectrometry for azithromycin (AZI), ciprofloxacin (CIP) and doxycycline (DOX). Angles between variables (arrows) reflect their correlations; arrow length is proportional to effect size. Samples (points) are colored by dehydration severity. All displayed variables have a significant effect (*P*<0.05, permutation test) except for ICP2, ICP3, and doxycycline (Table S4). For the RDA: *R^2^*=0.25 and adjusted *R*^2^=0.184, permutation test *P* = 0.001. To improve readability, collection date and location are not shown (see Fig. S6 for these details). Color code in all panels: blue: mild dehydration, orange: moderate, and red: severe.

We focused on ICP1 for subsequent analyses given its prevalence. The distribution of ICP1 relative abundance was variable and less clearly associated with dehydration status than *Vc* (**Fig. S3**). Deeper investigation revealed that it was not simply the presence of phage that mattered, but the ratio of ICP1 to *Vc*. Higher ratios were inversely associated with dehydration severity (**Fig. 1B**); the same results were obtained using qPCR rather than metagenomics to quantify the ratio (**Fig. S4**). This simple ratio is therefore a potential biomarker of ‘effective’ phage predation that could be used in clinical, diagnostic, and epidemiological applications.

As a preliminary proof of concept, we tested the hypothesis that the ICP1:*Vc* ratio could be used to delineate the dehydration status. We used a bootstrapping method to identify an optimal ratio to differentiate between patients with dehydration (moderate or severe) versus patients without dehydration by WHO clinical measures (Methods). This step is clinically important because patients with moderate and severe dehydration (‘positives’ in this analysis) require rehydration treatment. The analyses yielded a threshold ratio of 0.18 (approximately 1 ICP1 to 5 *Vc*). At this threshold, the approach had a sensitivity of 85% (95%CI 80% to 89%) and positive predictive value (PPV) of 95% (95%CI 92-96%) to identify dehydrated patients; the specificity was 53% (95%CI 24% to 72%) and negative predictive value was 26% (95%CI 19% to 35%). The samples were distributed as 248 true positives, 16 true negatives, 14 false positives, and 45 false negatives. Clinically, a high sensitivity is preferred over high specificity in order to not ‘miss’ dehydrated patients; the high PPV gives justification to expend resources for a fluid resuscitation. The results demonstrate the potential utility of the phage:bacteria ratio as a biomarker to differentiate severity status and requires an independent study for validation and further evaluation of the low NPV.

We next sought to identify potential interactions between ICP1 and temporal stages of disease. Previously, ICP1 was found to be associated with early, rather than late stages of disease, peaking on the first day of infection in cholera patients sampled over time (*36*). Given that we collected one sample per patient at hospital admission, we were unable to determine with certainty whether ICP1 suppresses *Vc* or whether it is a non-causal marker of late-stage disease when patients are recovering. Despite this limitation, we recorded self-reported duration of diarrhea, providing a proxy for disease progression. We found that higher relative abundances of ICP1 were associated with mild dehydration at early stages of disease (duration of diarrhea <72h) but not at later stages (**Fig. S3B and D**). We therefore favor a scenario in which ICP1 suppresses *Vc* at early disease stages in a way that reduces disease severity. However, further time series studies will be required to establish causality.

### Antibiotics in stool are inversely associated with disease severity

To visualize the complex relationships between disease severity, bacteria, and phages in the context of antibiotic exposure, we used redundancy analysis (RDA; **Fig. 1C; Table S4**). For simplicity, the seven most dominant bacterial species identified by principal component analysis were included (**Fig. S5**). As explanatory variables, we visualized clinical data with strong effects, chosen by forward selection and starting with phages and antibiotic concentrations (**Fig. S6**). In accordance with the indicator species analysis (**Fig. 1A, Table S3**), higher *Vc* relative abundance was positively correlated with severe dehydration (**Fig. 1C**). ICP1 was moderately associated with *Vc*, consistent with a phage’s reliance on its host for replication, but less associated with severe dehydration. The antibiotics azithromycin (AZI) and to a lesser extent ciprofloxacin (CIP) were negatively correlated with *Vc* and severe dehydration, suggesting their role in suppressing cholera infection and disease. Supporting previous reports that AZI suppresses *Vc* (*37*), AZI was most strongly anticorrelated with *Vc* in our cohort (**Fig. 1C**). We did not identify annotated antibiotic resistance genes associated with AZI exposure (**Fig. S7**) at established thresholds (**Table S5**. In contrast, CIP exposure was significantly associated with the resistance genes *dfrA* and *aph6* (**Fig. S8),** which are both associated with *Vc* in our metagenomes (**Fig. S9**) and have previously been linked with CIP resistance in *Vc* (*16, 38*). Taken together, these results suggest CIP exposure selects for resistance genes within patients, potentially explaining why CIP may be less effective at suppressing *Vc* than AZI.

### Azithromycin suppresses predator-prey dynamics

We next asked if and how antibiotics interact with phages to suppress *Vc* within patients. To do so, we modeled the relationships between ICP1, *Vc* and antibiotic exposure within each patient. We fit generalized additive models (GAMs) of *Vc* (relative abundance from metagenomics or absolute abundance from qPCR) as a function of ICP1, antibiotic concentrations, and their interaction, including dehydration status as a random effect. We fit GAMs with all quantified antibiotics and their interaction with ICP1, as well as separate models for each antibiotic, alone or in combination, and compared them based on their Akaike Information Criterion (AIC; **Tables S6, S7**). The most parsimonious model (with the lowest AIC), using either metagenomics or qPCR data, showed a significant negative relationship between *Vc* and AZI (**Fig. 2A, S10**). This result is consistent with the redundancy analysis (RDA) results (**Fig. 1C**) and with known patterns of *Vc* suppression by AZI during infection (*37*). The relationship between *Vc* and ICP1 was quadratic in both metagenomics-and qPCR-based models: at low ICP1 abundance, the relationship was positive but became negative at higher ICP1 abundance (**Fig. 2B, S10**). This alternation between positive and negative correlations is consistent with predator-prey dynamics within infected patients (*39*). However, at higher concentrations of AZI, the quadratic relationship flattened, effectively suppressing the phage-bacteria interaction, likely because AZI kept *Vc* at low density. In the future, these interactions could be interrogated using patient time-series and laboratory experiments challenging *Vc* with antibiotics and phages.

**Fig. 2.**
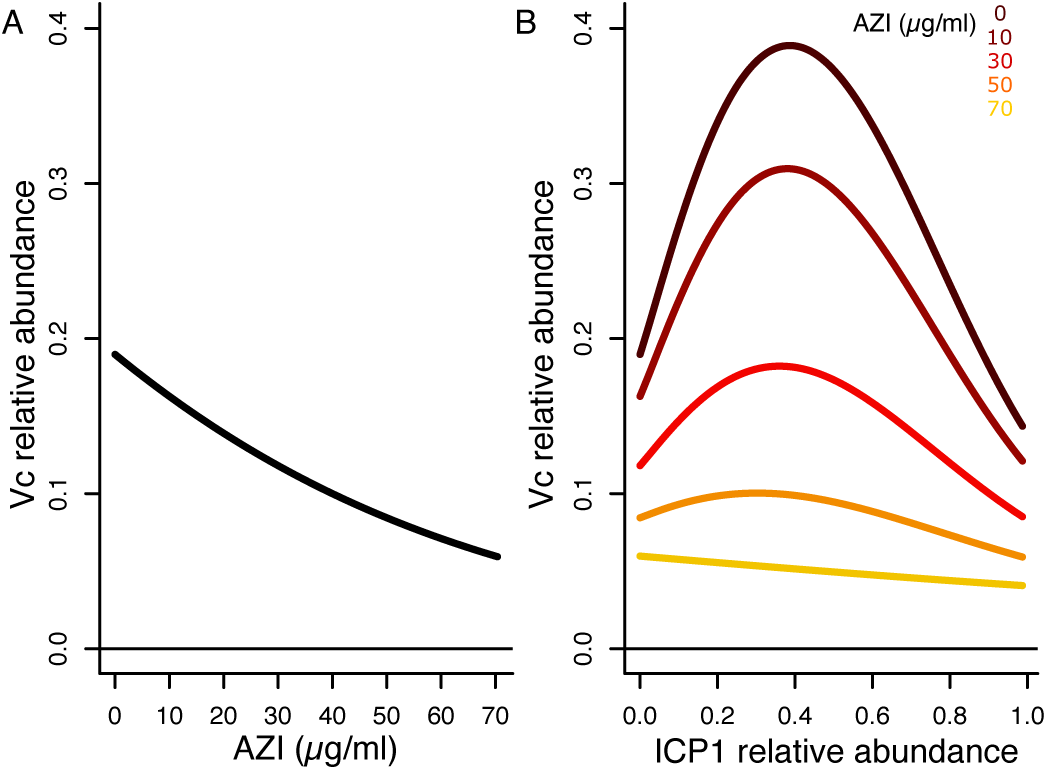
Interactions between *V. cholerae*, phage ICP1, and azithromycin. Generalized additive models (GAM) results, fit with relative abundance of *Vc* as a function of antibiotic concentrations (µg/ml) and ICP1 relative abundance in 344 metagenomes. (A) *Vc* declines in relative abundance with higher abundance of azithromycin (AZI) in µg/ml. (B) The relationship between ICP1 and *Vc* is affected by AZI concentration (µg/ml); the illustrated AZI concentrations show regular intervals between the minimum (0 µg/ml) and maximum (70 µg/ml) observed values. Both effects of AZI (A) and the ICP1-AZI interaction (B) are significant (Chi-square test, *P*<0.05). For details on GAM outputs see Table S6. Relative abundances are from metagenomics; see Fig. S10 for equivalent analyses using qPCR data.

### Integrative and conjugative elements (ICEs) are associated with phage suppression

The ICE is a region of the *Vc* genome that encodes resistance to both antibiotics and phages (*13*). ICEs have conserved ‘core’ genes along with variable ‘hotspots’ encoding different genes; for example, hotspot 5 is a ∼17kb region associated with phage resistance. At the time of our sampling, ICEVchInd5 (abbreviated here as *ind5*) and ICEVchInd6 (*ind6*) were the two most prevalent ICE types in Bangladesh (*17*). These ICEs differ in their anti-phage systems: *ind5* encodes a type 1 bacteriophage exclusion (BREX) system while *ind6* encodes several other different restriction-modification systems (*17*).

We screened for ICEs in metagenomes by mapping reads against reference sequences for *ind5* (NCBI accession GQ463142.1) and *ind6* (accession MK165649.1). An ICE was defined as present when 90% of its length was covered by at least one metagenomic read (**Fig. S11A**). We found that 64% (144/224) of samples with *Vc*>0.5% or ICP phages >0.1% of metagenomic reads contained *ind5*, 12% (26/224) contained *ind6*, and 24% (54/224) had no detectable ICE. The lack of ICE detections was not due to the lack of *Vc* in a metagenome because ICE-negative samples did not contain fewer *Vc* reads (**Fig. S11B**).

Resistance mechanisms on ICEs have been shown to suppress phage *in vitro* (*17*), but their relevance within human infection remains unclear. We found that metagenomes without a detectable ICE (denoted as ICE-) were associated with higher phage:*Vc* ratios (**Fig. 3**). The effect was strongest for ICP1, which had the largest sample size (**Fig. 3A**). This observation is consistent with ICE-encoded mechanisms suppressing phage within patients. Higher ICP1:*Vc* ratios, which occurred more often in ICE-patients, were also associated with mild dehydration (**Fig. 3B**). ICP1 is more strongly suppressed by *ind5* than by *ind6* (**Fig. 3**), while ICP3 appears to be better suppressed by *ind6* than *ind5*, albeit with borderline statistical significance **(Fig. S12**). We next compared ratios by phage resistance genotype (*ind5, ind6* vs ICE-) and dehydration status. For patients with mild dehydration, we observed lower ICP1:*Vc* ratios in *ind5* compared to ICE-samples (Kruskall-Wallis test with Dunn’s post-hoc correction, *p =* 0.022). Despite this difference, some patients with mild disease and *ind5* still had non-zero ICP1:*Vc* ratios (**Fig. 3B**), indicating the ICP1 is imperfectly suppressed by *ind5*. In the severe group, *ind5* patients also had lower ICP1:*Vc* ratios than ICE-patients (Dunn’s post-hoc test with BH correction, *P*=0.0035). In the moderate group, patients carrying *ind5* had lower ICP1:*Vc* ratios compared to patients with *ind6* (Dunn’s post-hoc test with BH correction, *P*=0.048), consistent with *ind5* more effectively suppressing ICP1. The same associations were evident using qPCR-based quantification of phage:*Vc* ratios (**Fig. S13**). Together, these results implicate ICEs in phage resistance during human infections, complementing and generally confirming the predictions of earlier laboratory experiments (*17*). That said, the suppression is not complete, and further experiments are needed to dissect the underlying causal relationships.

**Fig. 3.**
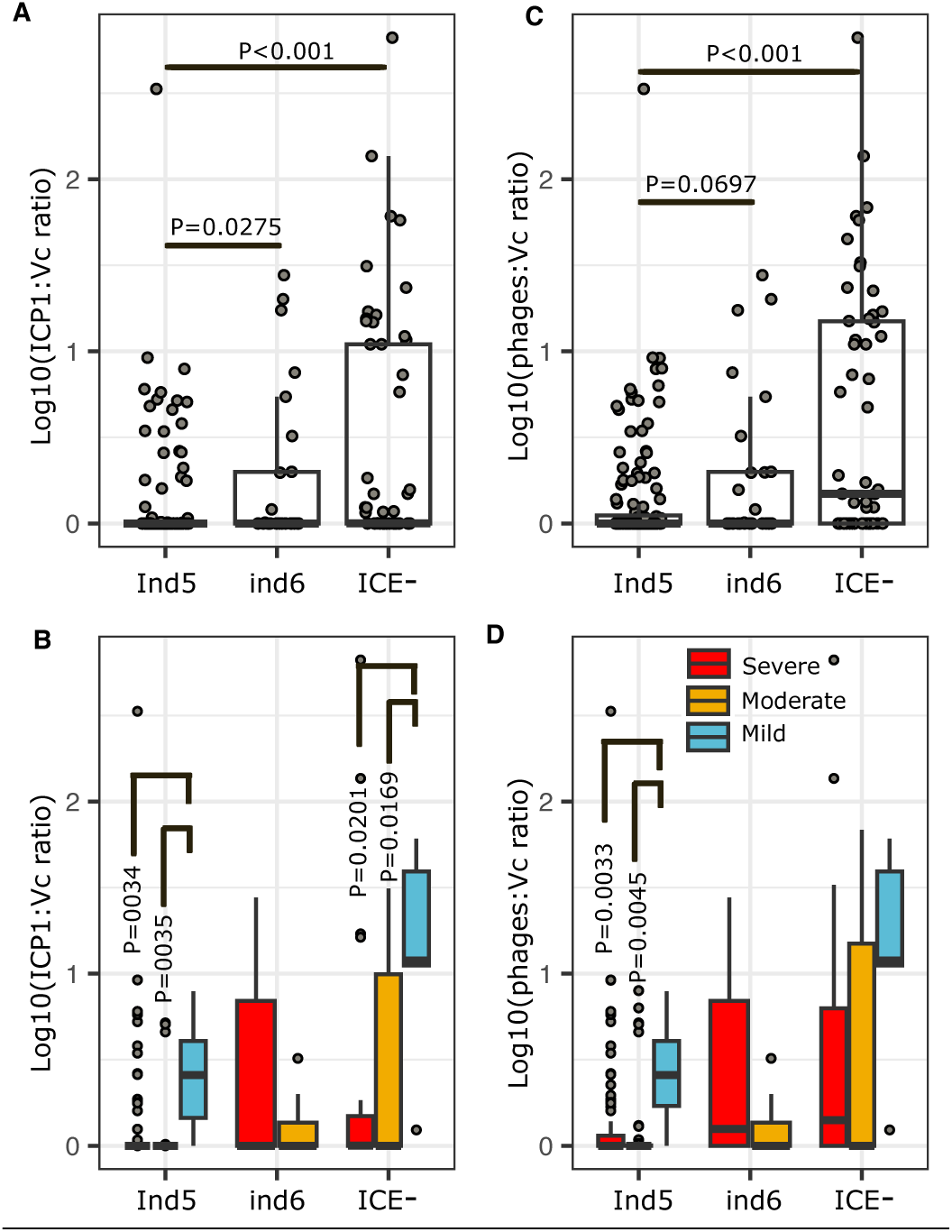
Integrative conjugative elements (ICEs) are associated with lower ICP1:*V. cholerae* ratios in patient metagenomes. (A) Distribution of ICP1:*Vc* ratios across patients with different ICE profiles. (B) The same data as (A) binned into boxplots according to dehydration status: mild (blue), moderate (orange) and severe (red). (C) Distribution of phage:*Vc* ratios, including the sum of all phages (ICP1, ICP2, ICP3). (D) The same data as (C) binned into boxplots according to dehydration status. *P*-values are from a Kruskal-Wallis test with Dunn’s post-hoc test adjusted with the Benjamini-Hochberg (BH) method. Only *P*-values < 0.1 are shown. Only samples with appreciable *Vc* or ICP1 were included (224 samples with *Vc*>0.5% or phages >0.1% of metagenomic reads), of which 54 samples were ICE-, 26 were *ind6*+ and 144 were *ind5*+. The Y-axes were log10 transformed after adding one to the ratios. The solid horizontal line is the median and the boxed area is the interquartile range. Relative abundances are from metagenomics; see Fig. S13 for supporting analyses using qPCR data.

### Hypermutation generates V. cholerae genetic diversity

In addition to variation in gene content in ICEs and other mobile elements, we hypothesized that resistance to phages and antibiotics could be conferred by point mutations (single nucleotide variants; SNVs) that existed before or emerged *de novo* during infection. Although we cannot fully exclude mixed infections by different *Vc* strains as a source of within-patient diversity, we found no evidence for more than one strain co-infecting a patient in our study population (**Fig. S14**). We previously found a low level of *Vc* genetic diversity within individual cholera patients (*40*) – on the order of 0-3 detectable SNVs – with the exception of hypermutation events characterized by DNA repair defects and dozens of SNVs in the *Vc* genome, primarily transversion mutations (*41*). Hypermutation generates deleterious mutations, but may also rapidly confer phage resistance, as shown experimentally with *Pseudomonas fluorescens* (*18*). Here, we were able to better quantify the frequency of hypermutators in a larger sample size, and test if within-host *Vc* diversity is associated with phage or antibiotic exposure – both of which could potentially select for resistance mutations to each factor.

To identify hypermutators in metagenomes, we tallied *Vc* populations with defects (nonsynonymous mutations) in any of 65 previously defined DNA repair genes (*42*) or with a relatively high number of SNVs (25 or more) (*41*). We used InStrain (*43*) to quantify *Vc* within-host diversity in 133 samples passing stringent sequencing quality filters (Methods) and found that 35% of samples (47/133) had both a high SNV count and nonsynonymous mutations in DNA repair genes – making them likely to contain hypermutators. Higher SNV counts were significantly associated with DNA repair defects (Fisher’s exact test, *P*<2.2e-16), consistent with these defects yielding higher mutation rates within patients. The number of SNVs was not confounded by *Vc* genome coverage (**Fig. S15A**). Consistent with our previous study (*41*), putative hypermutators had a distinct mutational profile enriched in transversions (**Fig. S15B,C**). For subsequent analysis, we considered all SNVs together, regardless of whether they were generated by hypermutation.

### Phages, not antibiotics, are associated with Vc within-host diversity

We hypothesized that *Vc* within-host diversity would be shaped by potential selective pressures, namely phages or antibiotics within patients. To test this hypothesis, we fit generalized linear mixed models (GLMMs) with phages and antibiotics as predictors of the number of high-frequency nonsynonymous (NS) SNVs in the *Vc* population within a patient. We focused on higher-frequency SNVs (>10% within a sample) as likely beneficial mutations (unlikely to rise to such high frequency by chance if neutral or deleterious) and on NS SNVs as more likely to have fitness effects. We fit several models with different combinations of predictors: a model with all antibiotics and their interaction with ICP1 and separate models with each antibiotic and its interaction with ICP1. We added *Vc* abundance as a fixed effect to the model to control for any coverage bias in SNV calling (**Table S8**). The most parsimonious model included *Vc* abundance and the interaction between *Vc* and ICP1 as predictors of the number of high-frequency NS SNVs. Adding antibiotics, or their interaction with ICP1, did not improve the model (**Table S8**), suggesting a limited role for antibiotics in selecting for *Vc* point mutations within patients.

In the model, *Vc* relative abundance and the interaction between *Vc* and ICP1 both had significant effects (GLMM, Wald test, *P*=0.00246 and *P*=0.00494 respectively). The negative relationship between *Vc* and the number of high-frequency NS SNVs (**Fig. 4A**) was not easily explained by sequencing coverage, since the total number of SNVs is not associated with *Vc* relative abundance (**Fig. S15A**). The number of high-frequency NS SNVs rose with increasing ICP1 – but only when *Vc* abundance was relatively high (**Fig. 4A**). As a control, we ran the same GLMM on NS SNVs without a minimum frequency cutoff and found no significant effects, suggesting that the interaction between ICP1 and *Vc* on SNV count is specific to high-frequency mutations, rather than low-frequency mutations that are more likely selectively neutral or deleterious. These data support a scenario in which ICP1 selects for NS SNVs when the *Vc* population is large enough to respond efficiently to selection – for example, at the beginning of an infection.

**Fig. 4.**
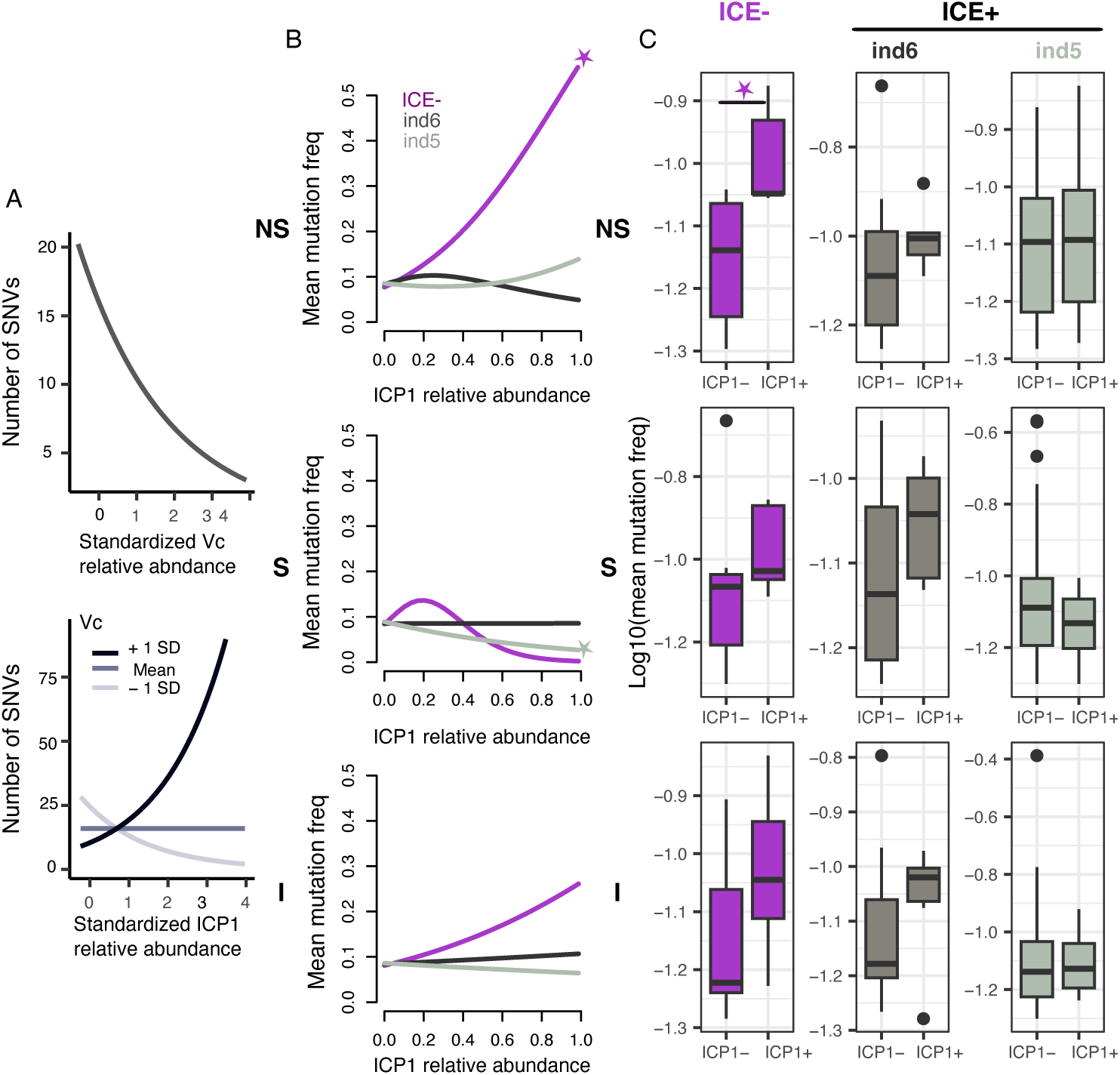
ICP1 selects for non-synonymous point mutations in the *V. cholerae* genome in the absence of ICE. (A) Results of a GLMM modeling high frequency nonsynonymous SNV counts as a function of *Vc* and ICP1 standardized relative abundances. In the bottom panel, shades of gray indicate *Vc* relative abundance at the mean or +/-1 standard deviation (SD) across samples. Both *Vc* and the interaction between *Vc* and ICP1 have significant effects (Wald test, *P*<0.05), the model was fit using 68 samples in which InStrain identified NS mutations at frequency > 10%. (B) GAM results with the mean mutation frequency as a function of the interaction between ICP1, ICE and mutation type (non-synonymous; NS, synonymous; S, or intergenic; I). Significant effects are shown with a star (Chi-square test, *P*<0.05). The model was fit using 130 samples that passed the post-InStrain filter for SNV quality (Methods). (C) Boxplots of mutation frequency in the presence or absence of ICP1 and/or ICEs. The only significant comparison is indicated with a star (Wilcoxon test, *P*=0.0094). Boxplots include 130 samples, of which 32 are ICP1+ (ICP1>=0.1% of reads) and 98 are ICP-(ICP1<0.1% of reads). The solid horizontal line is the median and the boxed area is the interquartile range. For supporting analysis using qPCR data, see Fig. S17.

If phages select for beneficial mutations, we expect these mutations to increase in frequency at higher intensity of phage predation. We lack time-series data from individual patients, but the relative abundance of phage provides a proxy for the combined effects of the strength and duration of phage selection. To test this hypothesis, we fit a GAM with the average within-sample frequency of SNVs as a function of ICP1, antibiotics, and their interactions. We included the fixed effect of ICE (*ind5*, *ind6,* or ICE-) as another factor that could provide phage or antibiotic resistance, as well as mutation type to differentiate among non-synonymous (NS), synonymous (S), and intergenic (I) mutations. We fit GAMs with all antibiotics and their interaction with ICP1, as well as models with or without each antibiotic separately (**Table S9**). The most parsimonious model included the interaction between ICP1, ICE and mutation type, but not antibiotics. ICP1 was a strong predictor of higher frequency NS SNVs in the absence of a detectable ICE (**Fig. 4B**). Samples in this analysis were unambiguous in terms of their ICE presence/absence patterns (**Fig. S16**).

To confirm and visualize this model prediction, we compared the distribution of the average frequency of NS SNVs between ICP1-positive and ICP1-negative samples. Consistent with the model, NS SNV frequency was significantly higher in ICP1-positive samples when the ICE was not detected (Wilcoxon test, *P*=0.0094) making this SNV category likely to contain targets of selection by ICP1 predation (**Fig. 4C**). Qualitatively similar results were found when ICP1 was quantified by qPCR instead of metagenomics (**Fig. S17**). Together, the results suggest that, in the absence of detectable ICE-encoded phage resistance, ICP1 selects for nonsynonymous point mutations instead. We identified several *Vc* genes, including some with membrane or virulence-related functions, that were mutated at higher ICP1:*Vc* ratios (**Table S10**); these provide candidate phage resistance mechanisms that can be explored in future experiments. In contrast, the secreted hemolysin gene, *hlyA,* that we previously observed to be mutated more often than expected within cholera patients (*41*) was among the genes most frequently mutated in patients with relatively low ICP1:*Vc* ratios (**Table S11**). This suggests that *hlyA* sequence evolution may be affected directly or indirectly by phage predation, through mechanisms that remain unclear.

As *Vc* evolves as a function of ICP1, we expect ICP1 evolution to be impacted by *Vc.* Specifically, we hypothesized that ICP1 may evolve to infect *Vc* in the presence of *ind5*, explaining some of the variation in both ICP1:*Vc* ratios and disease severity (**Fig. 3**). Despite the generally low genetic diversity of ICP1 (*21*), we were able to quantify SNVs in the ICP1 genome in 45 samples. This sample size was too low to fit sophisticated models, but simple correlations allowed us to draw tentative conclusions. First, we ruled out sequencing coverage as source of bias in SNV calling (Spearman correlation between ICP1 relative abundance and number of SNVs, *p* > 0.1 for all SNV categories). Next, we observed a negative correlation between the ICP1:*Vc* ratio and the number of NS SNVs in the ICP1 genome *–* a correlation that is only significant when *Vc* encodes an *ind5* ICE (**Fig. 5A, S18**). This is consistent with our observation that *ind5* is associated with lower ICP1:*Vc* ratios in our cohort (**Fig. 3**), potentially suppressing ICP1 and applying selection to escape suppression via NS mutations. Some of these NS mutations may be beneficial to the phage, rising to high frequency along with ICP1 – which is indeed what we observe: the mean frequency of NS SNVs in ICP1 increases with the ICP1:*Vc* ratio, but only in the presence of *ind5* (**Fig. 5B, S19**). Several ICP1 genes, mostly hypothetical proteins, repeatedly acquired NS mutations in the presence of *ind5*, providing candidate escape mutations to test in future work (**Table S12**).

**Fig. 5.**
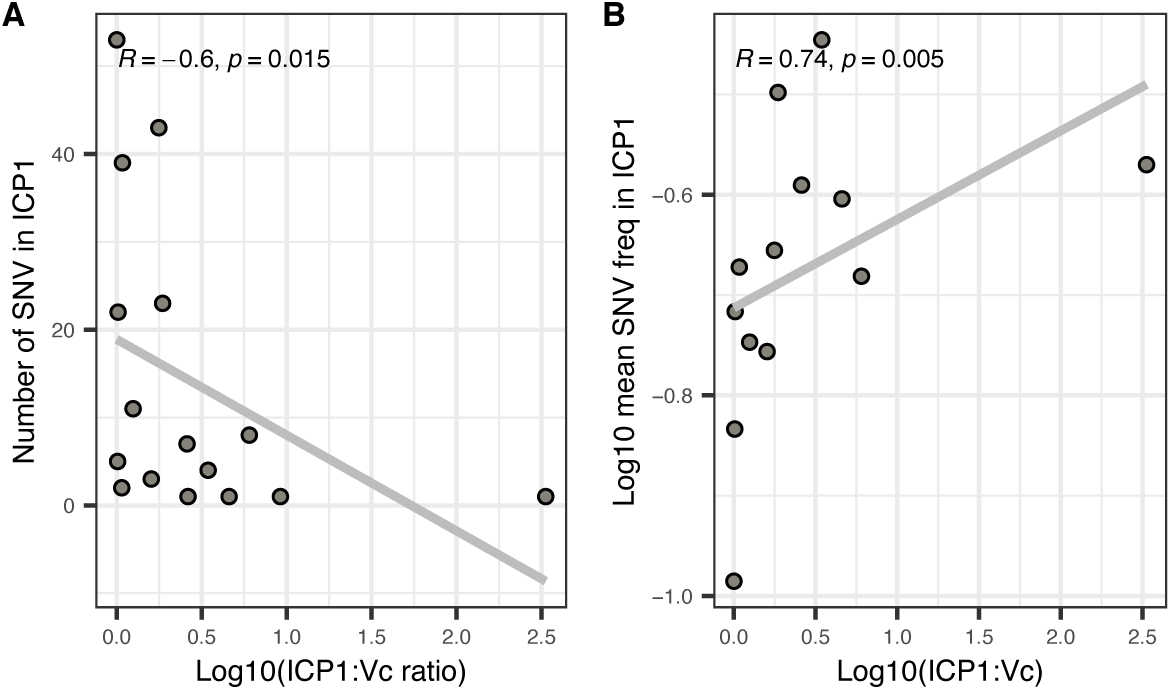
ICP1 evolution in samples containing ICE *ind5*. (A) The number of nonsynonymous (NS) SNVs detected in the ICP1 genome is negatively correlated with the ICP1:*Vc* ratio in the presence of *ind5*. (B) The mean frequency of NS SNVs in the ICP1 genome is positively correlated with the ICP1:*Vc* ratio in the presence of *ind5.* The X-axes were log10 transformed after adding one to the ratios. The Spearman correlation coefficients and *p*-values are shown. See Figures S18 and S19 for equivalent plots in ICE-and *ind6* samples, and for synonymous and intergenic SNVs.

## Discussion

The tripartite interactions between pathogens, phages, and antibiotics have been studied in the laboratory, *in silico* with mathematical models, and to a lesser extent in the field, but how these factors interact during human infection remains an open question. Our objective was to characterize these interactions in the context of cholera. We analyzed more than 300 stool metagenomes from cholera patients enrolled at hospital admission across Bangladesh during an entire outbreak season. We found that high predator (ICP1) to prey (*Vc*) ratios were inversely associated with disease severity and provide a proof of concept for translational applications. We demonstrated how *Vc* and ICP1 interact within patients, with ICP1 selecting for potential phage resistance point mutations in the absence of ICE-encoded anti-phage defenses, and *Vc* selecting for point mutations in the phage genome in the presence of *ind5*. This apparent coevolution between predator and prey likely has longer-term consequences for cholera infection and transmission. Antibiotics, particularly azithromycin, also played a role in suppressing *Vc* and could mask phage-bacteria interactions. Ciprofloxacin was associated with known antibiotic resistance genes, but we found no evidence that antibiotics select, as ICP1 does, for high-frequency nonsynonymous point mutations. Thus, although resistance mechanisms to certain phages and antibiotics colocalize to the ICE (*17*), they impose distinct selective pressures that could be exploited to improve the efficacy of antibiotics by combining them with phage therapy.

Our study has several limitations. First, samples were collected at a single time point at hospital admission which allowed us to establish statistical correlations, but we cannot infer causality in the absence of time-series or interventional clinical studies. Second, our cohort allowed us to study the interaction between *Vc* and ICP1, but the sample size for ICP2 and ICP3 was insufficient for most statistical analyses. Third, we prioritized common antibiotics for mass spectrometry, but we cannot exclude a role for other unmeasured antibiotics. Fourth, due to logistical limitations, we extracted DNA from a bacterial pellet plus a small amount of supernatant from each sample. This allowed us to capture both intracellular and cell-bound phages, along with free phage particles, but sequencing each fraction separately could yield further insights into distinct phage populations. From a clinical perspective, the measurement of dehydration status was categorical and could be improved in future studies by incorporating digital tools to quantify the degree of dehydration (*44–46*). Finally, our study lacked information about host genetics or immunity, which also influence disease severity (*8, 47*). Future studies combining rich patient metadata, time-series design, long-read metagenomics, and isolate genome sequencing will complement and build upon these findings.

## CONCLUSION

We propose that an index of effective phage predation, quantified as the phage:bacteria ratio, might be used as a tool for physicians to assess disease severity, and potentially prognosticate a disease course. We show here that this ratio is associated with cholera disease severity, but its predictive value should be studied in larger cohorts sampled over the course of infection. Whether this biomarker can be generalized to phages other than ICP1 and diseases other than cholera, and whether the association with disease severity changes as predator and prey coevolve over time, remains unclear. The potential of phage therapy and prophylaxis has long been recognized, and a combination of ICP1, 2, and 3 prevents cholera in animal models (*35*). However, just as hypermutators can drive the evolution of resistance to combinations of antibiotics (*48*), they may also help pathogen populations to survive combinations of phages, while increasing their potential to evolve resistance to future antimicrobial treatments. Phage therapy cocktails will therefore need to be updated regularly to remain effective against currently circulating coevolved bacteria, and creative new strategies are needed to minimize the unwanted evolution of phage, and possibly antibiotic, resistance.

## Materials and Methods Summary

### Ethics Statement

The samples analyzed were collected within two previously published IRB approved clinical studies in Bangladesh: (i) The mHealth Diarrhea Management (mHDM) cluster randomized controlled trial (IEDCR IRB/2017/10; icddr,b ERC/RRC PR-17036; University of Florida IRB 201601762) (*46*). (ii) The National Cholera Surveillance (NCS) study (icddr,b ERC/RRC PR-15127) (*49*); See supplementary materials for further details.

### Study Design

The study design was a prospective longitudinal study of patients presenting with diarrheal disease at five Bangladesh Ministry of Health and Family Welfare district hospitals (both mHDM and NCS sites) and two centralized NCS hospitals (BITID; icddr,b) from March 2018 to December 2018. Sites were distributed geographically nation-wide (*50*). See supplementary materials.

### Sample collection

Stool samples were collected at hospital admission. Aliquots for transport and subsequent culture were stabbed into Cary-Blair transport media; aliquots for molecular analysis were preserved in RNAlater (Invitrogen). See supplementary materials.

### Microbiological and molecular analysis

Culture was performed via standard methods (*51*); total nucleic acid (tNA) was extracted from the RNAlater preserved samples using standard methods. Criteria for subset subsequent shotgun metagenomic sequencing were: (i) culture positivity, (ii) phage (ICP1,2,3) detection by PCR among culture-negative samples, or (iii) *Vc* detection by PCR among a random 10% of culture-negative and phage (ICP1,2,3) negative samples. Sequencing libraries were prepared using the NEB Ultra II shotgun kit and sequenced on Illumina NovaSeq 6000 S4, pooling 96 samples per lane, yielding a mean of >30 million paired-end 150bp reads per sample. Among samples identified for metagenomic analysis, qPCR was performed to determine absolute abundances of *Vc*, ICP1, ICP2, and ICP3.

### Antibiotic detection by liquid chromatography mass spectrometry (LC-MS/MS)

Those cholera samples identified for metagenomic analysis were analyzed by qualitative and quantitative LC-MS/MS for antibiotics. Based on prior research (*16, 37*), the target list prioritized 5 common antibiotics: ciprofloxacin, doxycycline/tetracycline, and azithromycin were analyzed quantitatively, and metronidazole and nalidixic acid were analyzed qualitatively. Standard curves were made for each quantitative target by preparing a dilution series of the three native and isotopic forms of the quantitative targets; clinical samples were spiked with the isotopes of the quantitative targets as internal references. See supplementary materials.

### Metagenomic data analysis

We taxonomically classified short reads using Kraken2 (*52*) and Bracken v.2.5 (*53*). Reads were assembled using MEGAHIT v.1.2.9 (*50*) and binned with DAS tool (*54*). We inferred probable phage contigs using geNomad v1.7.0 (*55*) and predicted their likely bacterial hosts with iPhoP v1.3.1 (*56*). To characterize intra-patient *Vc* diversity, we used StrainGE (*57*) and InStrain v.1.5.7 (*43*). To identify antibiotic resistance genes in metagenomes, we used deepARG v 1.0.2 (*58*). See supplementary materials for details.

### Statistical analyses

Statistics and visualizations were done in R version 3.6.3 and R studio version 1.2.5042. See supplementary materials for details.

## Competing interests

Authors declare that they have no competing interests.

## Data and materials availability

All sequencing data are deposited in the NCBI SRA under BioProject PRJNA976726. See supplementary materials for further information.

## Code availability

Computer code needed to reproduce figures and results in this study is available on Github at https://github.com/Naima16/Cholera-phage-antibiotics. DOI: 10.5281/zenodo.10573867 (77).

## Acknowledgements

We thank the patients for participating in this study as well as the clinical and laboratory teams who collected the samples. We are grateful to S. Flora and colleagues at the Institute of Epidemiology, Disease Control and Research (IEDCR), Ministry of Health and Family Welfare, Government of Bangladesh who collaborated on the original clinical studies in which the samples analyzed herein were collected. We are also grateful to R. Autrey, B. Johnson and K. Berquist for their administrative expertise at the University of Florida. AIK and FQ were the principal investigators (PI) in Bangladesh and PIs of the ERC/RRC approvals at the icddr,b. EJN was the PI and obtained IRB approval at the University of Florida. Associated protocol numbers/registrations are: IEDCR IRB/2017/10; icddr,b ERC/RRC PR-17036; University of Florida IRB 201601762; clinicaltrials.gov NCT03154229. This collective research infrastructure and support was invaluable to the success of this study. We thank members of the Nelson, Khan, Qadri and Shapiro labs, and Frédérique Le Roux, for discussions that improved the manuscript.

## Funding

This work was supported by the National Institutes of Health grants to EJN [R21TW010182] and KBB [S10 OD021758-01A1] and internal support from the Emerging Pathogens Institute at the University of Florida and the Departments of Pediatrics/ Children’s Miracle Network (Florida). BJS and NM were supported by a Canadian Institutes for Health Research Project Grant. AC was supported by a Postdoctoral Mobility Fellowship of the Swiss National Science Foundation [P500PB_214356]. The funders had no role in study design, data collection and analysis, decision to publish, or preparation of the manuscript.

## Author contributions

Conceptualization: FW, AIK, BJS, EJN

Methodology: NM, MAS, ETC, MTRB, YB, ACM, MK, AC, EF, AV, LSB, KBS, BJS, EJN

Investigation: NM, MAS, ETC, KI, MIUK, YB, ACM, MK, AC, EF, AV, LSB, BJS, EJN

Visualization: NM, AC, BJS, EJN

Funding acquisition: KBS, FW, AIK, BJS, EJN

Project administration: MTRB, YB, KBS, AIK, BJS, EJN

Supervision: KBS, FQ, AIK, BJS, EJN

Validation: NM

Formal analysis: NM, BJS, EJN

Resources: BJS, AIK, EJ

Data Curation: NM, EJN, AIK

Writing – original draft: NM, ETC, MAS, ACM, MK, LSB, KSB, AIK, BJS, EJN

Writing – review & editing: NM, AIK, AC, BJS, EJN

## Supplementary materials

Materials and methods

Figures: S1 to S19

Tables: S1 to S12

Data Files S1 to S5

References 59-77

## Notes

### Competing Interest Statement

The authors have declared no competing interest.

### Summary of Updates

- added cutpointr analysis as preliminary proof of concept of the diagnostic potential of the ICP1:V. cholerae ratio - minor edits for clarity

