## Supplementary Methods, Figures, and Tables for "Phage predation, disease severity and pathogen genetic diversity in cholera patients"

**Affiliations:**
[1] Department of Microbiology & Immunology, McGill University, Montréal, Canada
[2] McGill Genome Centre, McGill University, Montréal, Canada
[3] Departments of Pediatrics and Environmental and Global Health, University of Florida, Gainesville, FL, USA
[4] Infectious Diseases Division (IDD) & Nutrition and Clinical Services Division (NCSD), International Centre for Diarrhoeal Disease Research, Bangladesh (icddr,b), Dhaka, Bangladesh
[5] Department of Chemistry, University of Florida, Gainesville, FL, USA
[6] McGill Centre for Microbiome Research, McGill University, Montréal, Canada

[*] Corresponding authors

**The PDF file includes:**
Table of contents
Materials and methods
Results
Figures: S1 to S19
Tables: S1 to S12
References 59-77

**Data files:**

**File S1.** Patient metadata.

**File S2.** Antibiotic concentrations (μg/ml) in stool samples.

**File S3.** Full list of *Vc* genes containing mutations in samples where ICP1 > *Vc* (% of reads).

**File S4.** Full list of *Vc* genes containing mutations in samples where ICP1 < *Vc* (% of reads).

**File S5.** Full list of ICP1 genes containing mutations in samples containing ICE *ind5*.

### MATERIALS AND METHODS

**Ethics Statement.**

The samples analyzed were collected within the context of two previously published IRB approved clinical studies conducted in Bangladesh: the mHealth Diarrhea Management (mHDM) cluster randomized controlled trial (IEDCR IRB/2017/10; icddr,b ERC/RRC PR-17036; University of Florida IRB 201601762; clinicaltrials.gov NCT03154229) (*46*) and a hospital-based National Cholera Surveillance (NCS) study (icddr,b ERC/RRC PR-15127) (*49*); the consent processes are described within these published studies.

**Study Design.**

The study design was a national prospective longitudinal study of patients presenting with diarrheal disease at five Bangladesh Ministry of Health and Family Welfare district hospitals (both mHDM and NCS sites) and two centralized hospitals (NCS sites alone) (BITID, icddr,b) from March 2018 to December 2018. Recruitment was based on the census of diarrheal disease patients that sought care at the hospitals. For the mHDM study, inclusion criteria were patients two-months of age or older, and presentation with acute diarrhea defined as three or more episodes of loose stools in the 24 hours prior to admission and a duration of disease less than seven days. For the NCS study, patients with diarrheal disease of all ages were included. For those under 2 months of age, diarrhea was defined as a change in stool habit from 'usual' (increased frequency with less formed stool). For those two months and older, diarrhea was defined as three or more loose or liquid stools within 24 hours or three loose/liquid stools or fewer causing dehydration in the last 24 hours. There was no exclusion based on enrollment in a second study (e.g., mHDM vs NCS). Patient metadata is in Data File S1.

**Sample Collection.**

For the mHDM study, the intent was to collect four stool samples per study site per day. For the NCS study, the intent was to collect daily stool samples from two participants less than five years old and from two participants five years of age and older; if the target number for an age group was not met, samples were collected from the other age group to achieve a total of four samples per day. For the samples collected at the five district hospitals and BITID, an aliquot was placed in transport media (Cary-Blair) and transferred to the icddr,b laboratory for culture, and a 0.5 ml aliquot was added to 1.3 ml of RNAlater to stabilize the sample for subsequent analysis; cold-chain below 4°C was not consistently available at these sites. For samples collected at the icddr,b, samples were stabilized in RNAlater; culture was performed directly. Samples were stored at the centralized icddr,b laboratory at -80°C.

**Microbial and molecular analyses.**

Standard methods were used to extract total nucleic acid (tNA) from the samples stabilized in RNAlater (Invitrogen). In brief, the samples were thawed at room temperature and centrifuged for five minutes at max speed. All but 50 µl of the supernatant was removed; the supernatant was refrozen for subsequent mass spectrometry. The remaining pellet, with the 50 µl supernatant, was combined with 380 mg of glass beads and 1 ml InhibitEx buffer and processed using the QIAamp Fast DNA Stool Mini Kit. Elutions were performed with 50 µl of Qiagen ATE buffer followed by a second elution with 50 µl ATE, quantification (Nanodrop) and storage at -80°C. For qPCR, we used established *tcpA* primers for *V. cholerae* (*37, 59*) and 16S ribosomal gene (rDNA) primers as a positive control for sample processing and extraction (*60*). Primer pairs were developed for the virulent bacteriophages ICP1 (gp58.2), ICP2 (gp24), and ICP3 (gp19) (**Table S1**); primer sequences are conserved between Southeast Asia and African isolates. Reactions used Luna qPCR master mix (New England Biolabs) with 25 µl volumes, and the method was a two-step reaction with denaturation at 98°C and annealing/extension at 60°C for all primers except ICP2 GP24 F/R (annealing/extension 63°C). Assays were performed on an Azure Cielo Real-time qPCR System and two Bio-Rad CFX96 Touch Real-Time PCR Detection Systems. Samples were analyzed in 96-well formats with two replicates except for 16SrDNA (one replicate); a Ct threshold less than 28 was used to define positivity (*37, 59, 61*). For quality control, samples that were negative for 16S rDNA were excluded from subsequent analyses. A third replicate was performed when: both replicates were positive but the difference in Ct values was more than 4, or one replicate was positive and one was negative and the difference in Ct values was greater than four. After the third replicate was performed, outliers were removed and the Ct values were averaged and the final determination of positivity was scored (Ct<28). Primers were validated using standard curves of (i) synthetic templates and (ii) biologic agents for *V. cholerae*, ICP1, ICP2, ICP3. Conventional methods were used for PCR.

**Antibiotic detection by liquid chromatography mass spectrometry (LC-MS/MS).**

*LC-MS/MS methodology for both qualitative and quantitative approaches***.** The experimental approach used in this study was based on prior studies (*16, 37*). Stool supernatants from the primary collection were obtained by centrifugation without filtration to minimize loss. Proteins were precipitated (1:7 ratio, volume:volumbe, of water:methanol).

Supernatants were diluted with methanol and water (1:1 v/v) in 1% formic acid for liquid chromatography, and 5 µl of supernatant was injected for analysis. LC-MS/MS was performed on a 2.1 x 150-mm Hypersil Gold aQ column (particle size, 3 µm) using a high-performance liquid chromatography system (Thermo UltiMate 3000 series) with an LTQ XL ion trap mass spectrometer (Thermo Fisher Scientific). Mobile phases were 1% formic acid in water (A) and 1% formic acid in methanol (B) and held at a constant 5%B for 2 minutes before ramping to 95%B at 15 minutes where it was held for an additional minute before returning to starting conditions for a total run time of 25 min. Blanks were run in between every two samples, as well as before and after quality control samples and standards.

Eluent was ionized using electrospray ionization (ESI) in positive mode at a spray voltage of 5 kV, a nitrogen sheath gas flow rate of 8 L $min^{-1}$, and capillary temperature of 300°C. Two scan events were programmed to perform an initial scan from *m/z* 100 to 1000, which was followed by targeted collision induced dissociation based on a retention time and mass list. Retention time windows ranged from 1.7 min to 5.1 min, depending on the elution range of the standards at high and low concentrations. Masses were targeted for the most abundant adduct or ion associated with each antibiotic (typically the $[M+H]^+$ ion) with a *m/z* 1 window.

*LC-MS/MS methodology specific to qualitative analysis.* The target list for the qualitative analysis was narrowed from prior work that assayed for antibiotics and non-antibiotics (*16, 37*): acetaminophen, metronidazole, ondansetron, furazolidone, nalidixic acid, sulfamethoxazole, trimethoprim, omeprazole, ciprofloxacin, cephalexin, penicillin V, amoxicillin, doxycycline or tetracycline, ceftriaxone, erythromycin, and azithromycin. Based on this prior work, we narrowed our analysis to the common clinically relevant agents of ciprofloxacin, azithromycin, doxycycline/tetracycline (same mass), nalidixic acid and metronidazole. Qualitative data analysis was performed using extracted ion chromatograms and MSMS matching with a control antibiotic MSMS library using Xcalibur 2.2 SP 1.48 (Thermo Fisher Scientific).

*LC-MS/MS methodology specific to quantitative analysis.* The target list for the quantitative analysis was furthered narrowed to: ciprofloxacin, azithromycin, and tetracycline/doxycycline. A standard curve was made for each quantitative target by preparing a dilution series of a mix of the three native forms of the quantitative targets (ciprofloxacin, doxycycline, azithromycin). The dilutions were: 0.5, 0.25, 0.125, 0.063, 0.05, 0.02, 0.01 µg/ml. All samples, including standards, were spiked with a 0.25 µM mix of isotope labeled control antibiotics (ciprofloxacin d-8 (Millipore-Sigma), doxycycline d-5 (Isosciences), azithromycin d-5 (LGC standards).

A calibration curve was generated by plotting the ratio of the area under the curve for analyte peaks in the ion chromatograms to the area under the curve of the isotope peaks against known concentrations of the seven standards. The linear line of best fit produced by this plot was generated and used to extrapolate the quantity of drug in mHDM samples. The area under the curve of chromatogram peaks produced by the target compound was compared to that of the peak produced by the isotope spike in to generate a ratio. This ratio was then converted to a concentration (in units of µg/ml) using the calibration curve equation. Data analysis was performed manually by viewing the extracted ion chromatograms and MSMS from each sample and matching with an in-house antibiotic library using 2.2 SP 1.48 (Thermo Fisher Scientific). The detection of a drug was confirmed when a peak in the chromatogram at the proper retention time window was identified and the most abundant adduct or ion associated with that drug (typically the $[M+H]^+$ ion) with a m/z 1 window. Measured antibiotic concentrations are in Data File S2.

**Metagenomic data analyses.**

*Short read classification using kraken2/bracken.* We classified all short-read data with a Kraken2 database (*52*) containing bacterial, archaeal, and viral domains, along with the human genome and a collection of known vectors (UniVec_Core) from NCBI, in July 2020. A Bracken database (*53*) was also built with a read length of 150 bp and the default k-mer length of 35. Kraken2 and Bracken version 2.5 were run with default parameters.

*Phage identification from metagenomic assemblies.* We characterized phages in 224 metagenomes with either >0.5% *Vc* or >0.1% ICP phage relative abundances. We used geNomad (v1.7.0, min-score 0.7, max-fdr 0.1) (*56*) to identify (pro)-phages from 83,928 contigs identified as 'viral'. iPhoP (v1.3.1, min-score 90%) (*56*) was used to predict the bacterial hosts of the identified phage contigs, resulting in 627 contigs classified as '*Vibrio* phages'. We used two different approaches to further characterize the predicted '*Vibrio* phage' sequences: (i) BLASTn to compare these sequences against known *Vibrio* phages in the INPHARED (*62*) database and reference sequences of the TLC (*63*) and PLE satellite phages (*22*) (% identity >= 80%, query coverage >= 90%), and (ii) vContact2 (*64*) to build similarity networks including the putative *Vibrio* phages and genomes from the INPHARED NCBI RefSeq databases. We used clusters in the vContact2 network to harmonize the naming of the INPHARED database genomes (eg. all INPHARED genomes which clustered together with the ICP2 reference genomes, were called 'ICP2').

*Quality filtering.* Before calling SNVs within the *Vc* population, metagenomes were decontaminated of human and PhiX reads by mapping the reads against the GRCh 38 assembly of the human genome and the PhiX genomes, with bowtie2 version 2.3.5 (*65*).

Unmapped reads were used for subsequent analyses. Then, we used iu_filter_quality_minoche from the Illumina Utils package (v2.10) with default parameters (*66*) to trim adapters, remove duplicates and for quality filtering on the short reads.

*SNV profiling.* Short reads were assembled with MEGAHIT (*50*) version 1.2.9 with the default parameters (k-mers length: k=21 et maxk=99 and min contig length of 1000. The contigs were grouped into bins with concoct (v1.0.0) (*67*) and MetaBAT 2 (v2.12.1) (*68*) with default parameters. Bins were then aggregated together using DAS Tool version 1.1.0 (*54*). Then we used drep (*69*) version 2.0.0 with S_ani 0.98, to de-replicate the bins and identify unique metagenome-assembled genomes (MAGs) with a secondary clustering threshold of 98% and minimum completeness of 75% and maximum contamination of 25%. To prevent miss mapping of reads from other species to the *V. cholerae* genome, we competitively mapped reads from all samples with sufficient *V. cholerae* or phages (*Vc*>0.5% of reads or phages>0.1% of metagenomic reads) against the concatenation of the 677 unique MAGs and the ICP1 genome from NCBI (reference NC_015157.1, accessed 2021/12/01), using bowtie2 v 2.3.5 (*65*). To characterize within-patient *Vc* and ICP1 genetic diversity, we profiled the resulting bam files using inStrain v1.5.7 (*43*) with default parameters (minimum coverage of 5 to call a variant). We also ran gene-level profiling with inStrain using genes annotated with prodigal. We used prodigal v2.6.3 (*70*) with the default parameters to predict genes in the *Vc* MAG and ICP1 and annotated them with eggNOG-Mapper v2 with default parameters and eggNOG database v2 downloaded on April 2021 (*71*).

To infer potential mixed infections, we used strainGST from the strainGE toolkit (*57*). To do so, we compiled a reference database of publicly available *V. cholerae* assemblies, including (i) all complete genomes from NCBI RefSeq (n=106, accessed 2023/05/12) and (ii) all assemblies published on NCBI Genbank between 2015/01/01 and 2023/05/12 and originating from samples collected between 2015/01/01 and 2019/12/31 (n=758).

To reduce false positive SNVs, we applied a stringent post-InStrain filter: all positions with coverage <20x were removed. We removed the first and last 100 positions of every scaffold, as well as positions with coverage below 0.3*median and above 3*median (median coverage across all the positions in the sample). We also removed sites that did not pass the coverage filter in more than two samples. Finally, we removed 420 sites that were variable in a *Vc* isolate genome sequenced alongside the metagenomes. These sites were considered prone to sequencing error since they varied in an isolate genome that theoretically should contain no variation. After applying all these quality filters, we were left with 133 samples in which SNV calls could be called.

*Hypermutator definition.* Hypermutators were identified as samples with one or more nonsynonymous mutations in DNA repair genes (DNA repair defects) in *V. cholerae*, and with 25 or more SNVs, as defined previously (*42*).

*Antibiotic resistance gene identification.* Antibiotic resistance genes (ARGs) were predicted from short reads using deepARG (*58*) v 1.0.2 with default parameters. Deeparg is a deep learning model that can predict ARGs from short-read metagenomic data; it uses DeepARG-DB, a merged database from 3 databases: Antibiotic Resistance Genes Database [ARDB], Comprehensive Antibiotic Resistance Database [CARD], and UniProt. This database was downloaded in December 2021. We used the relative abundance of ARGs normalized by the 16S rRNA gene content in the sample. We assessed the relative abundances of 634 known ARGs conferring resistance to 34 antibiotic classes. To identify associations between these ARGs and gut microbes, we ran a multiple factor analysis (MFA) with the 37 most dominant species from a PCA (20 highest coordinates on both axes) and all the ARGs.

*SXT ICE identification.* We used Bowtie2/2.3.5 with the 'very-sensitive' option to map short reads against the two most prevalent SXT ICEs in Bangladesh at the time of our sampling (*17*): ICEVchind5 (NCBI accession GQ463142.1) and ICEVchind6 (NCBI accession MK165649.1). The ICE was considered as present in a metagenome when 90% of its length was covered by at least one read. Using this criterion, 144 samples (59%) were ind5+, 26 (10.6%) ind6+, and 54 (22.1%) ICE-.

**Statistical analyses.**

All statistics and visualizations were done in R version 3.6.3 and R studio version 1.2.5042; the MedCalc Software Ltd. Diagnostic test evaluation was used for sensitivity and specificity calculations (Version 22.019; accessed February 7, 2024). To reduce dimensions of the species composition table, we ran a principal component analysis (PCA) on the Hellinger transformed abundance data (5719 species*344 samples) (decostand function and rda function in the vegan R library) and selected the 20 most dominant species on each axis (37 in total) in the analyses, except for the redundancy analysis (RDA) where only the seven most dominant species were included for simplicity.

To compare ICP1:*Vc* ratios among different dehydration profiles, we performed a Kruskal-Wallis test (kruskal.test function from the stats R library) with Dunn's post-hoc test adjusted for multiple tests using the Benjamini-Hochberg (BH) method (dunnTest function from the FSA R library). To estimate the performance of the ICP1:*Vc* ratio as a biomarker of disease severity, we used the cutpointr function from the cutpointr R library. This function utilizes a bootstrapping method (fixed to 1000) to identify the optimal ICP1:*Vc* ratio that differentiates between mild and moderate or severe status. The selected cutpoint maximizes the aggregate values of sensitivity and specificity.

To understand relationships between taxa in the microbiome, phages, antibiotics, and patient metadata, we used an RDA on bacterial species abundances (rda function from the vegan R library). Only the seven most dominant species identified with the PCA were included: *Vibrio cholerae, Escherichia coli*, *Bacteroides vulgatus*, *Bifidobacterium longum*, *Shigella flexneri*, *Bifidobacterium breve*, and *Streptococcus mitis*. As explanatory variables, we used three antibiotics: azithromycin (AZI), ciprofloxacin (CIP), Doxycycline (DOX) as well as the three phages (ICP1, ICP2, ICP3), along with the most contributing patient metadata, selected with forward selection method (ordiR2step function from the vegan R library). These metadata are: the area where the sample was collected, date of sampling, age in years, vomiting state (yes or no) and dehydration status (severe, moderate, or mild). We began the forward selection with a model with phages and antibiotic concentrations (µg/ml). The RDA was run on log-chord transformed abundances, and a permutation test with 999 iterations was used to assess the statistical significance of both the model and of each explanatory variable (anova.cca function from vegan library in R). The explained variation $R^2$ and adjusted $R^2$ were estimated with the RsquareAdj function in the vegan R library.

To identify species associated with each degree of dehydration, we used the indicator species analysis (*72*) as implemented in the multipatt R function from the indicspecies R library (with no group combination, duleg parameter set to TRUE and 9999 permutations) on the log-chord transformed species table (334 samples, 37 dominant species from the PCA) using the decostand and rda functions from the R vegan library). Reported P-values correspond to a permutation test with 999 iterations.

To study associations between ARGs and species, we ran a multiple factor analysis (MFA) (*73*) using the mfa function from the FactoMineR R package, with the 37 most dominant species and all 634 resistance genes identified by deepARG. The visualization was done with the fviz_mfa_var function from the factoextra R library.

To model the relationships between phages, antibiotics, and *Vc* within each patient, we fit a generalized additive model (GAM) with *Vc* relative abundance as a function of ICP1, antibiotics, and the interaction between them. We added the degree of dehydration as a random effect because it improved the model (smallest Akaike information criteria (AIC) compared to the other models). We fit several GAMs with different combinations of predictors: from a model with all antibiotics and their interaction with ICP1 to separate models with each antibiotic and its interaction with ICP1 and compared them based on their AIC (AICtab function from the bblme R package). The most parsimonious model was retained (the one with the smallest AIC). The GAMs were fitted using a beta error distribution with log-link function because *Vc* relative abundance is a continuous value between 0 and 1. We evaluated the selected model fit by inspecting residual distributions and fitted-observed value plots using the gam.check function from the mgcv R library. All P-values reported for the GAMs correspond to the Chi-square test from gam.summary function from mgcv R package. We also reported the adjusted $R^2$ (from the same function) as an evaluation of the goodness of fit (**Table S6**). To support the results from the models fitted on metagenomic relative abundances, we repeated the same methods with the qPCR-based absolute abundances of *Vc* and ICP1. Since the response (*Vc* absolute abundance from qPCR) is a continuous variable, we used additive models (Gaussian GAMs) after log-transforming the response (**Figure S10**, **Table S7**).

We tested whether phages or antibiotics select for potentially adaptive mutations in *Vc* by fitting generalized linear mixed models (GLMMs) (glmmtmb function from glmmTMB R package) with phages and antibiotics as predictors of the number of high-frequency nonsynonymous SNVs in the *Vc* genome. We added *Vc* abundance as a fixed effect to the model to control for any coverage effects. We focused on higher-frequency SNVs (>10% within a sample) as more likely to be beneficial, and nonsynonymous SNVs as those more likely to have fitness effects. We fit several models with different combinations of predictors: from a model with all antibiotics and their interaction with ICP1 to separate models with each antibiotic and its interaction with ICP1 and compared them based on their AIC (AICtab function from bblme R package). The most parsimonious model was retained (the one with the smallest AIC). Because the response is count data, we fit three different count GLMMs: Poisson, negative binomial 1 and negative binomial 2, implemented in the glmmTMB R package, and then selected the model with the lowest AIC as described (*74*). The selected model was fitted with the nbinom2 error distribution. All continuous predictors were standardized to zero mean and unit variance before analyses to improve the model's convergence. We evaluated the selected model fits by inspecting the residuals using the DHARMa library in R (simulateResiduals and plot functions). All the *P*-values reported for the GLMMs correspond to the Wald *P*-values reported by the glmm.summary function from the R package glmmTMB. As an evaluation of the goodness of fit, we compared the GLMMs and the corresponding null models (the same model but with no fixed effects other than the intercept) with likelihood-ratio tests (anova function from the stats R package) and reported the corresponding P-values (**Table S8**). We also reported the marginal $R^2$ as a measure of the variance explained by fixed effects, estimated with the r2 function from the performance R package. We used GLMMs because they are currently more flexible than GAMs in the range of count models that they can fit (https://bbolker.github.io/mixedmodels-misc/glmmFAQ.html). Furthermore, they can deal with overdispersion with two versions of negative binomial distributions: negative binomial 1 and negative binomial 2, respectively with linear and quadratic parametrization (*75*).

To further test the hypothesis that phages or antibiotics select for adaptive mutations in *Vc*, we examined the relationship of the frequency of nonsynonymous SNVs in *Vc* with phages and antibiotics. We fit a generalized additive model (GAM) (gam function in mgcv R library) with the average frequency of nonsynonymous SNVs as a function of ICP1, antibiotics, and their interactions, and *Vc* and ICP1 relative abundances from metagenomic data. We also included the fixed effect of ICE presence/absence as another factor that could provide phage or antibiotic resistance as well as mutation type to differentiate among nonsynonymous, synonymous, and intergenic mutations. We fit GAMs with all antibiotics and their interaction with ICP1, as well as several simpler models with each antibiotic separately, and compared them based on their AIC. The GAMs were fitted using a beta error distribution with a log-link function because the average minor allele frequency of nonsynonymous SNVs is a continuous value between 0 and 0.5. We evaluated the selected model fit by inspecting residual distributions and fitted-observed value plots using the gam.check function from the mgcv R package. All the P-values reported for the GAMs correspond to the Chi-square test from the gam.summary function of the mgcv R package. We also reported the adjusted R2 (from the same function) as an evaluation of the goodness of fit (**Table S9**).

### SUPPLEMENTARY FIGURES

### FIGURE S1

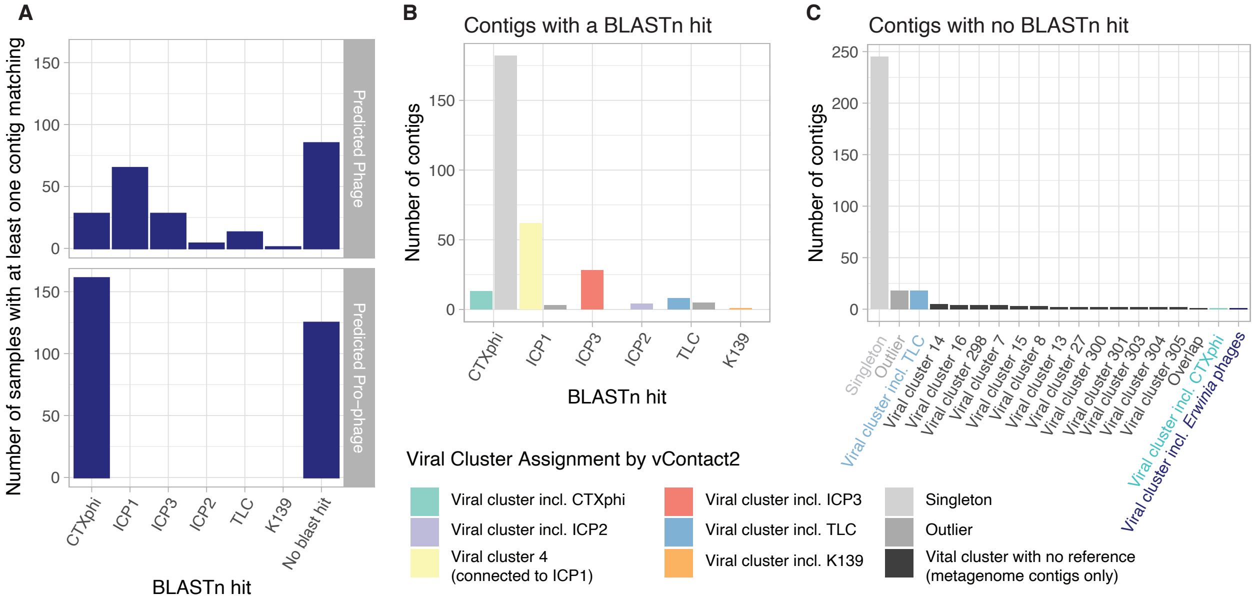

**Fig S1: Inferred *Vibrio* phages in 224 metagenome assemblies.** (A) At least one predicted *Vibrio* phage contig matched a CTXphi, ICP1, ICP3, TLC, ICP2 and K139 genome using BLASTn in 189, 63, 28, 13, 4 and 1 samples, respectively. (B) Phages ICP3, ICP2 and K139 are unambiguously assigned using vContact2. Contigs matching the CTXphi reference genome using BLASTn clustered either with the CTXphi reference viral cluster or formed singletons (observed in only one sample). Similarly, most contigs matching an ICP1 reference genome using BLASTn were assigned by vContact2 to Viral Cluster 4, which did not include any reference phage genomes, but numerous connections to the reference ICP1 genome. (C) For 321 inferred *Vibrio* phage contigs from 210 samples, there was no BLASTn hit to a known *Vibrio* phage. Of these, 245 formed singletons, 18 were labeled as ‘outliers’ and one was placed between two viral sequence clusters. The remaining 57 contigs with no BLASTn hit clustered into 16 viral clusters, 14 of which exclusively comprised ‘*Vibrio* phages’ identified in this analysis (mean cluster size = 2.8 contigs, range 2-5). Eighteen contigs with no BLASTn hit clustered with the TLC satellite phage (which flanks the CTXphi prophage in the *V. cholerae* genome), and one each with CTXphi and in a cluster predominantly including *Erwinia* phages.

**FIGURE S2**

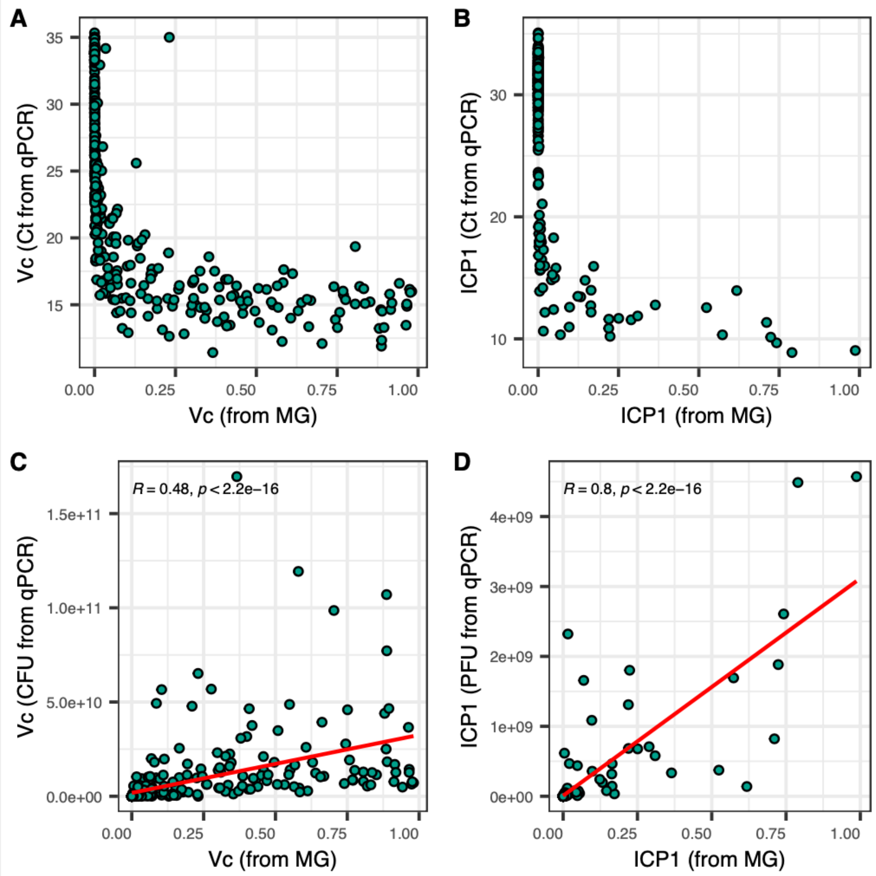

**Fig S2. Correlations between qPCR- and metagenomics-based quantification of *Vc* and ICP1.** Raw qPCR cycle threshold (Ct) values plotted against relative abundances in metagenomes for (A) *Vc* and (B) ICP1. Inferred number of (C) *Vc* colony forming units (CFU) per ml of stool and (D) ICP1 plaque forming units (PFU) per ml, both at a Ct threshold of 28. Red lines show Pearson correlations, with coefficients and *p*-values noted in the figure panels.

**FIGURE S3**

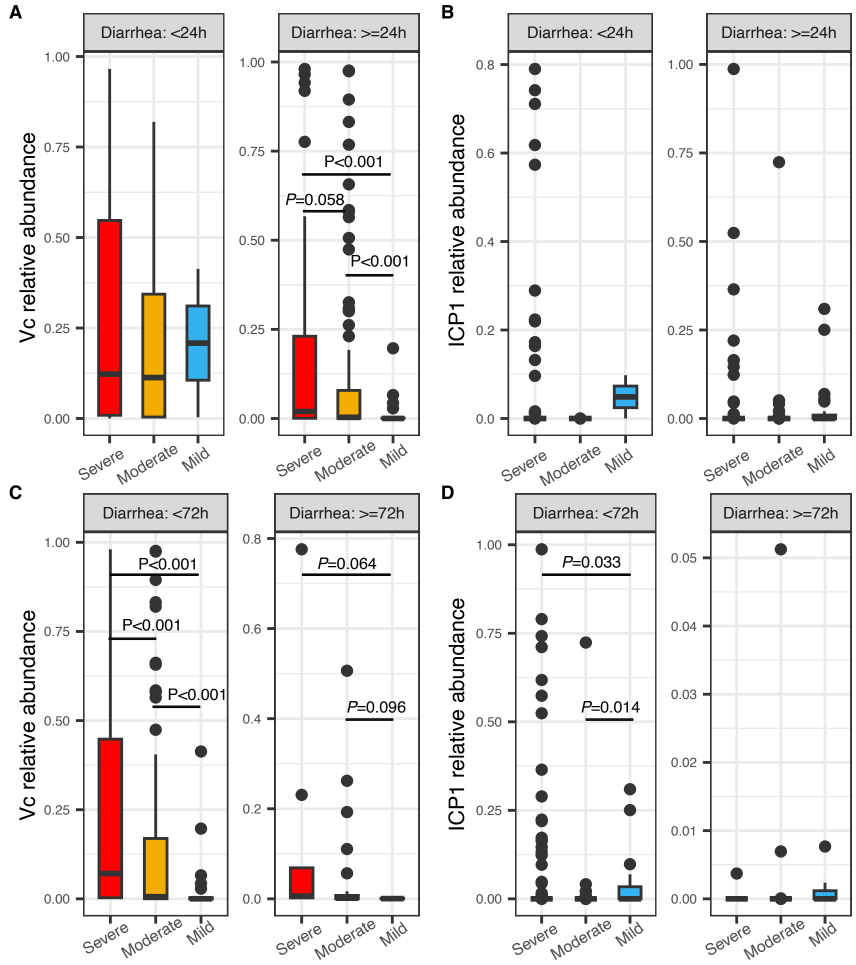

**Fig S3. ICP1:*V. cholerae* ratios among patients with different dehydration status binned by self-reported duration of diarrhea.** Panels show the relative abundance of *Vc* across dehydration status when duration of diarrhea is (A) <24h vs. >=24h, or (C) <72h vs. >=72h. ICP1 relative abundance is shown acros dehydration status when duration of diarrhea is (B) <24h vs. >24h, or (D) <72h vs. >=72h. *P*-values are from a Kruskal-Wallis test with Dunn's post-hoc test, adjusted for multiple tests using the Benjamini-Hochberg (BH) method. Only significant (*P*<0.05) and marginally significant *P*-values (<0.1) are shown. 323 samples with *Vc* >0% of metagenomic reads were included, with 165 from severe, 128 from moderate, and 30 from mild cases. The solid horizontal line is the median and the boxed area is the interquartile range.

**FIGURE S4**

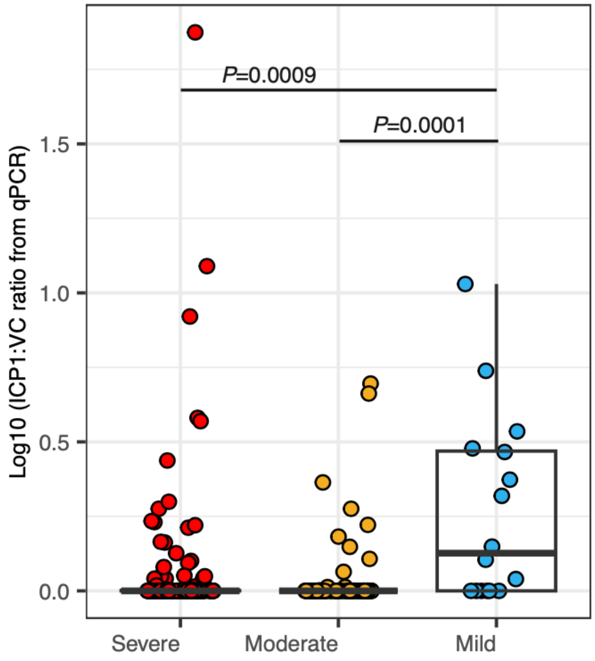

**Fig S4. The ICP1:*V. cholerae* ratio quantified with qPCR is higher in patients with mild dehydration.** *P*-values are from a Kruskal-Wallis test with Dunn's post-hoc test, adjusted for multiple tests using the Benjamini-Hochberg (BH) method. Only significant *P*-values (<0.05) are shown. Only 248 samples with *Vc*>0% from qPCR data were included to exclude zeroes from the denominator of the ratio, with 138 from severe, 94 from moderate, and 16 from mild cases. A pseudocount of one was added to the ratio before log transformation. This figure is a version of Figure 1B using qPCR data.

**FIGURE S5**

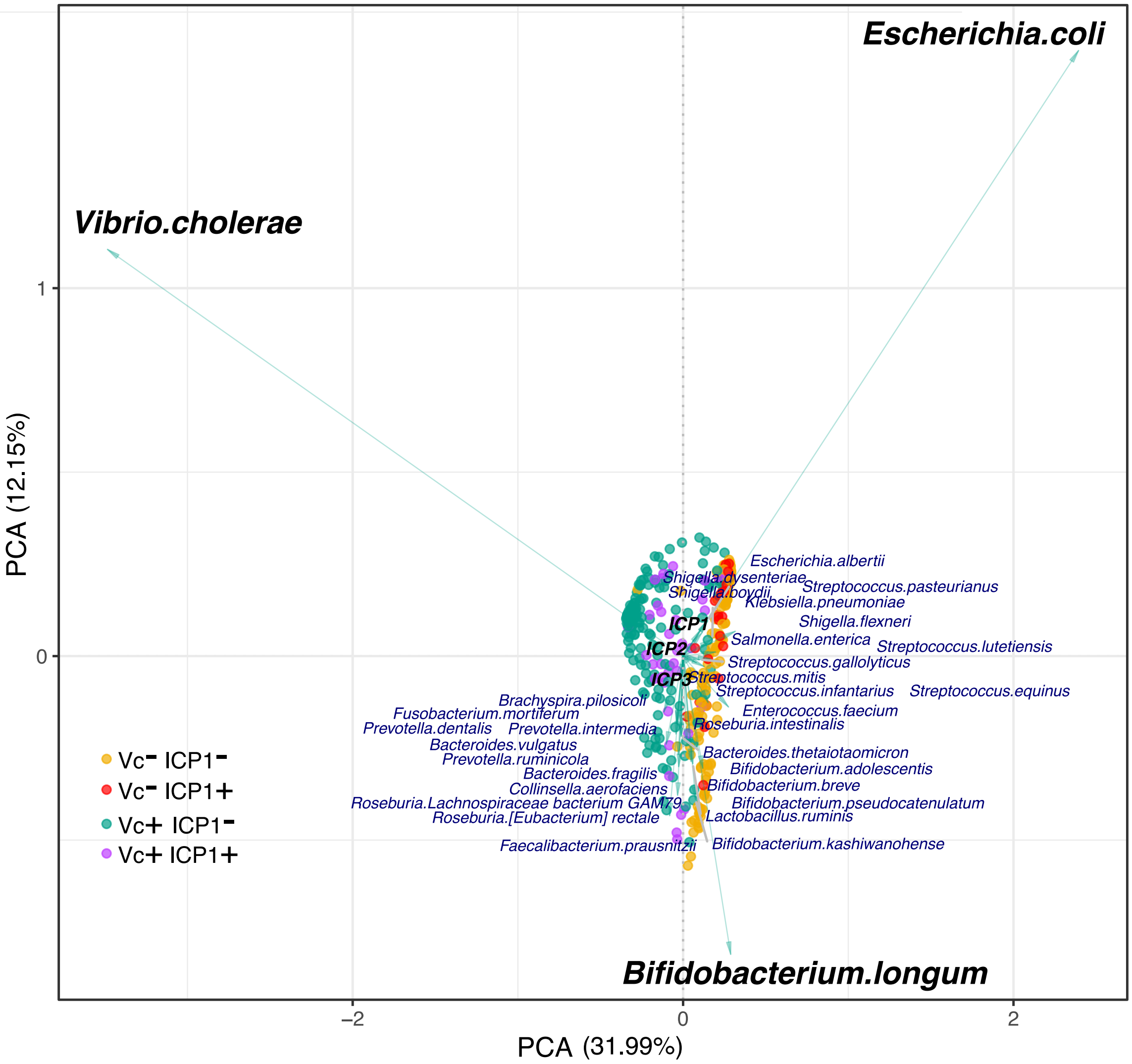

**Fig. S5. Principal component analysis (PCA) of the taxonomic composition of cholera patient stool samples.** This analysis was run to reduce species composition dimensions from 5719 annotated with kraken2/bracken to the 37 most dominant (with the 20 most dominant on the two first axes). Circles indicate samples; arrows indicate species. Samples are colored by their *Vc* and ICP1 relative abundances (percentage of metagenomic reads). Yellow: patients with *Vc* < 0.5% and ICP1 < 0.1%, red: *Vc* < 0.5% and ICP1 >= 0.1%, green: *Vc* >= 0.5% and ICP1 < 0.1%, purple: *Vc* >= 0.5% and ICP1 >= 0.1%.

**FIGURE S6**

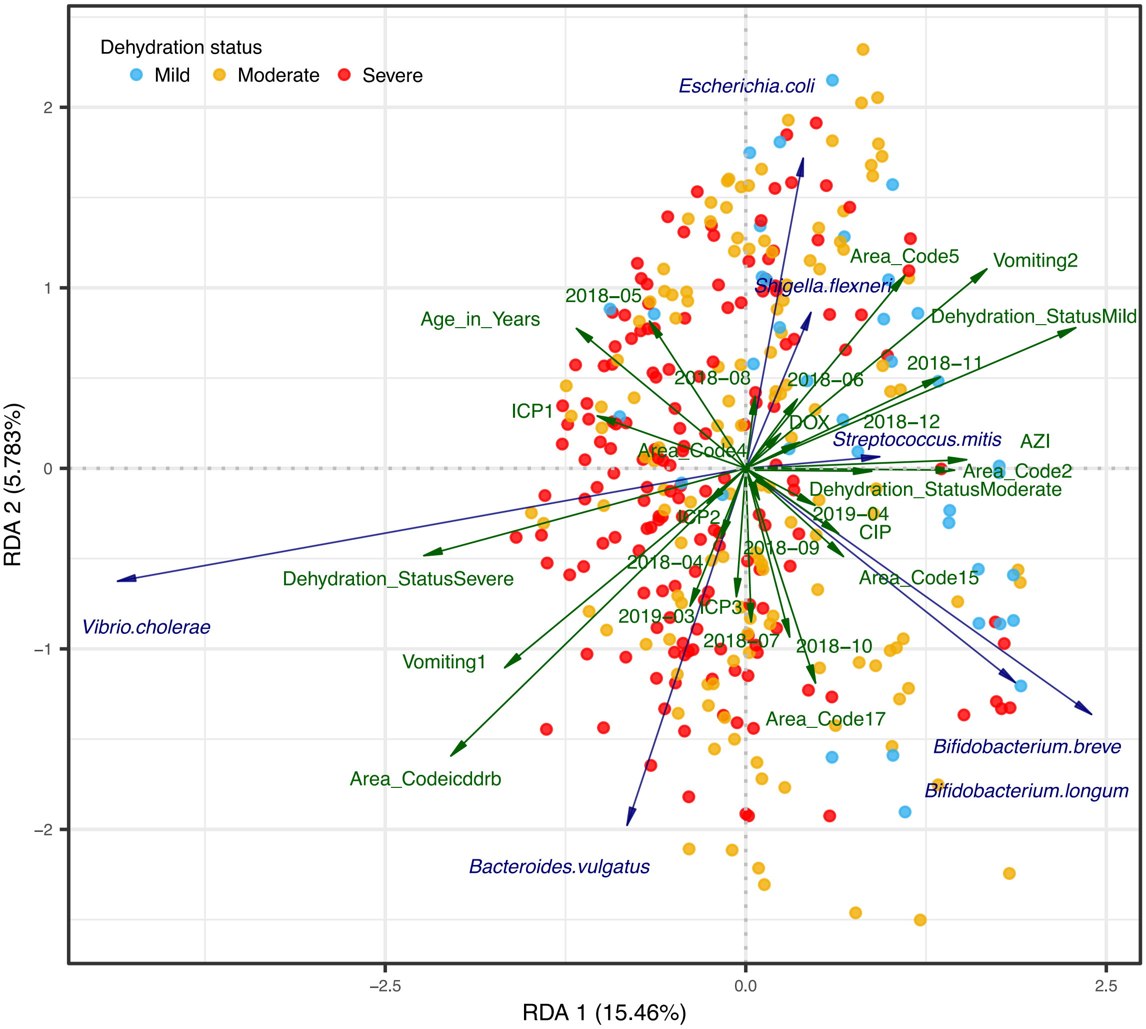

**Fig S6**. **Redundancy analysis (RDA) showing relationships among the seven most dominant bacterial species identified with PCA (Figure S5) and explanatory variables**: phages (ICP1, ICP2, ICP3), patient metadata (age in years, vomiting state (yes or no), dehydration status (severe, moderate or mild), the area where the sample was collected, and date of sampling), and antibiotic concentration from quantitative mass spectrometry for azithromycin (AZI), ciprofloxacin (CIP) and Doxycycline (DOX). Angles between variables (arrows) reflect their correlations; arrow length is proportional to effect size. Samples (points) are colored by dehydration severity: blue (mild dehydration), orange (moderate) and red (severe). All displayed variables have a significant effect ($P<0.05$, permutation test) except for ICP2, ICP3, and doxycycline (Table S4). For the RDA: $R^2$=0.25 and adjusted $R^2$=0.184, permutation test $P$ = 0.001. All variables are shown here; the collection date and area code were omitted in Fig. 1C for clarity.

**FIGURE S7**

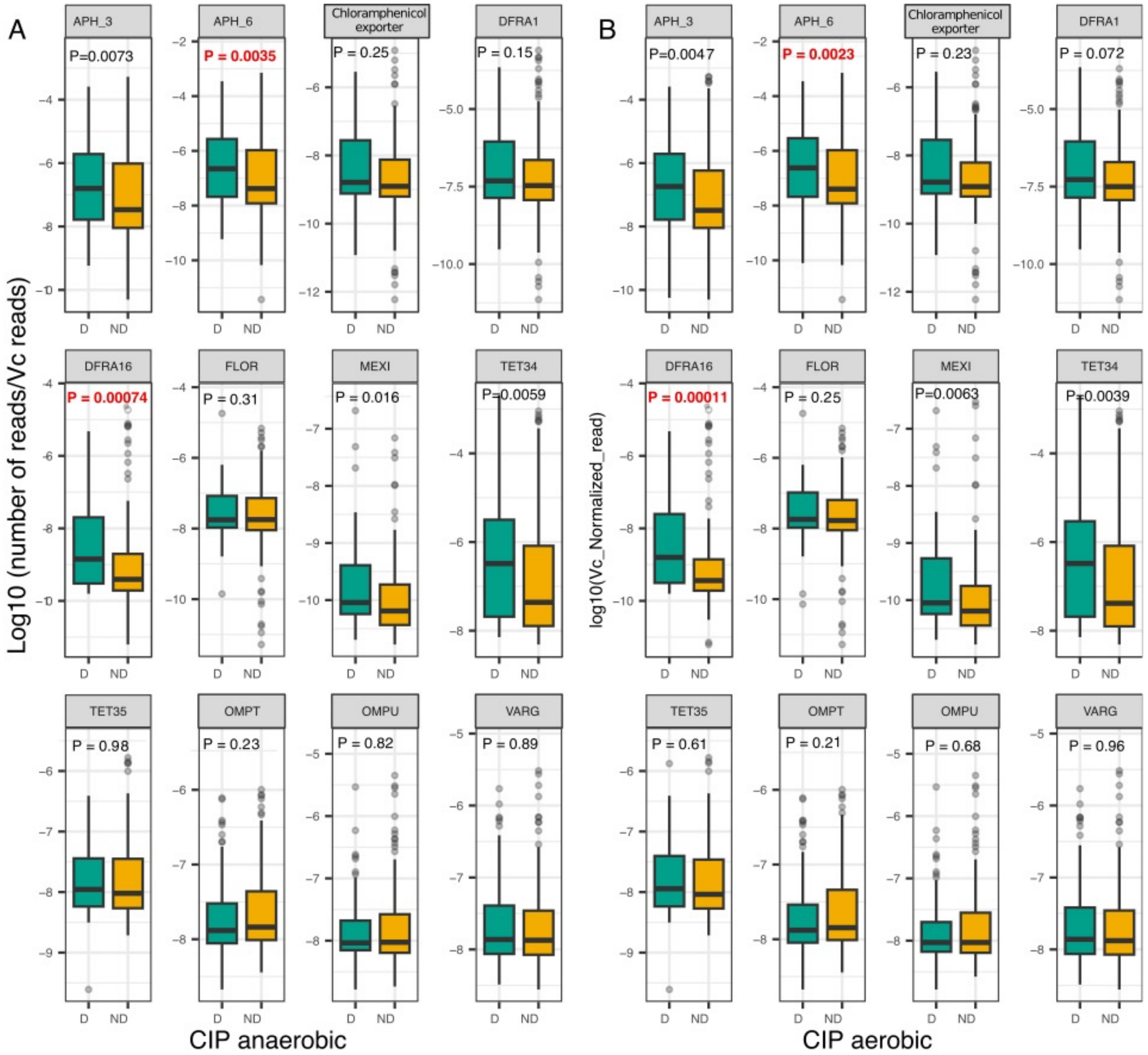

**Fig. S7. Distribution of the relative abundance of *V. cholerae*-associated ARGs in patients with different concentrations of ciprofloxacin (CIP).** CIP>=MIC (detected; D) and CIP<MIC (not detected; ND) under anaerobic conditions (A) or aerobic conditions (B). CIP MIC=0.016 µg/ml under aerobic conditions and 0.063 µg/ml under anaerobic conditions. The Y-axis is the relative abundance of ARGs in metagenomes normalized by 16S rRNA gene reads and by *Vc* reads. Only *Vc*-positive samples are plotted (*Vc*>0 reads, 323 out of 344 samples). *P*-values are from a Wilcoxon test. In red: BH corrected *P*<0.05 after correction for 24 tests.

**FIGURE S8**

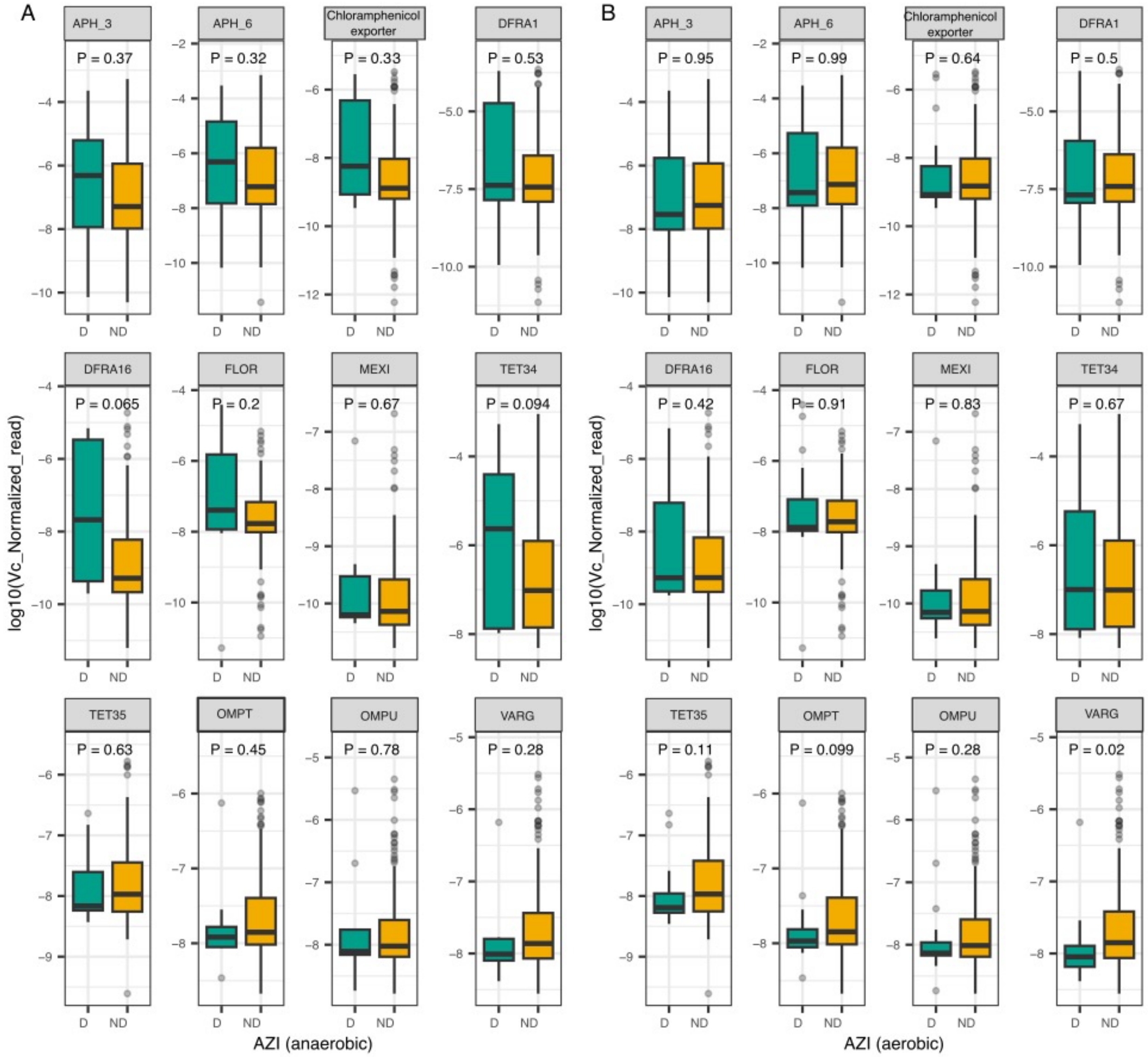

**Fig. S8. Distribution of the relative abundance of *V. cholerae*-associated ARGs in patients with different exposures to azithromycin (AZI).** AZI>=MIC (detected; D) and AZI<MIC (not detected; ND) under anaerobic conditions (A) or aerobic conditions (B). AZI MIC=1 μg/ml under aerobic conditions and 8 μg/ml under anaerobic conditions. The Y-axis is the relative abundance of ARGs in metagenomes normalized by 16S rRNA gene reads and by *V. cholerae* reads. Only *V. cholerae*-positive samples are plotted (*V. cholerae*>0 reads, 323 out of 344 samples). P-values are from a Wilcoxon test. No comparisons (D vs. ND) are significant (*P*<0.05) after BH correction for 24 tests.

**FIGURE S9**

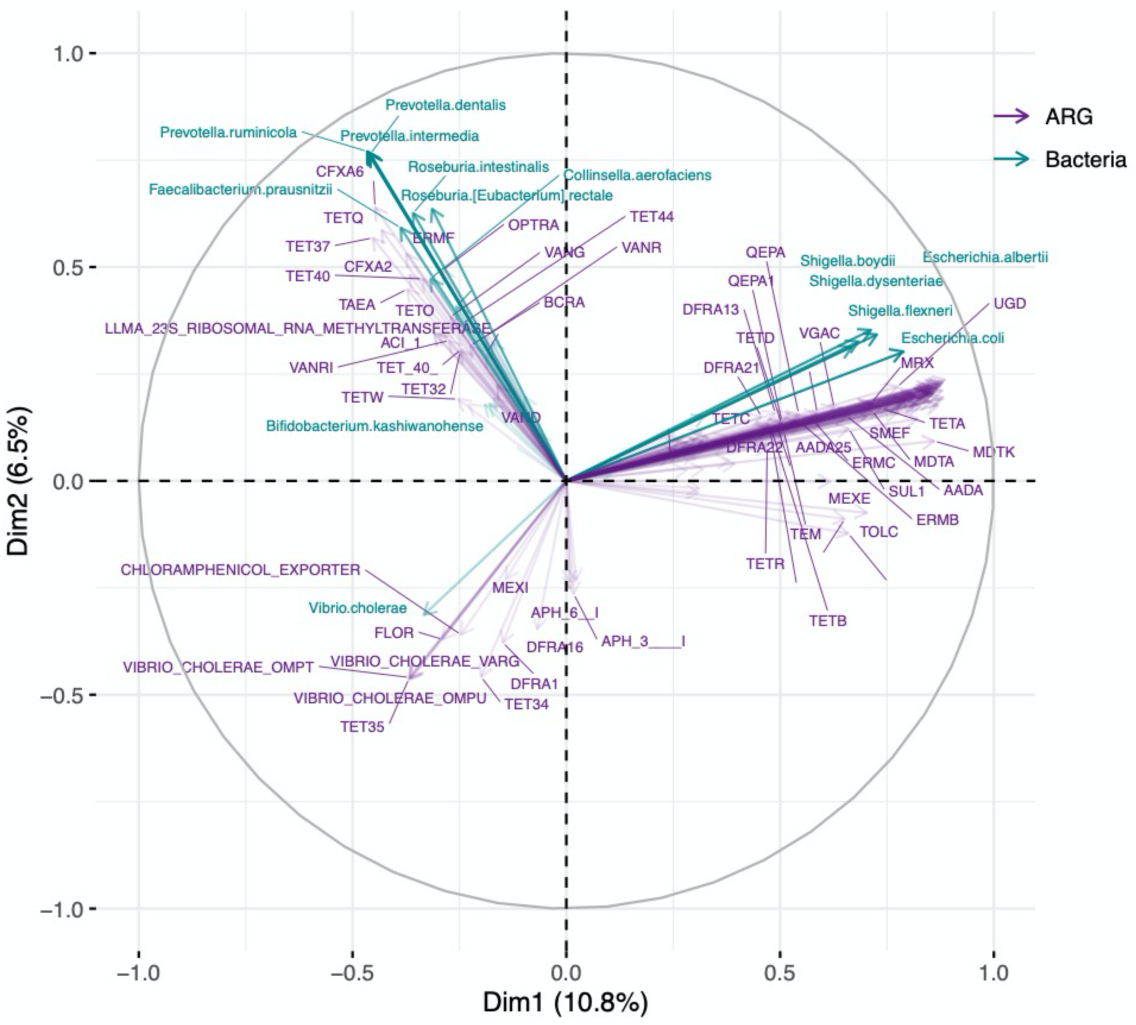

**Fig S9. Multifactor analysis (MFA) showing correlations between the relative abundance of bacterial species and the relative abundance of antimicrobial resistance genes (ARGs) in metagenomes normalized by 16S rRNA gene reads in patient gut microbiomes.** The 37 most dominant species from the PCA (Fig S5) and all the resistance genes annotated with deepARG were included in the ordination. 12 ARGs clustered with *Vc*: OMPU, OMPT, VARG, TET34, TET35, DFRA1, DFRA16, FLOR, Chloramphenicol exporter, Aph6, Aph3, MEXI. Only the greatest correlations on each axis are shown for clarity.

**FIGURE S10**

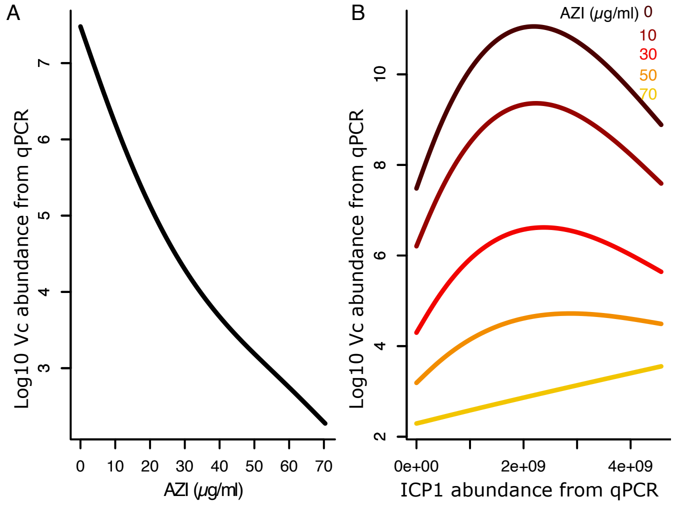

**Fig S10. Interactions between *V. cholerae*, phage ICP1, and azithromycin tested with qPCR data**. (A) *Vc* declines in absolute abundance with higher relative abundance of azithromycin (AZI) in µg/ml. (B) The relationship between ICP1 and *Vc* abundances estimated with qPCR is affected by AZI concentration (µg/ml); the illustrated AZI concentrations show regular intervals between the minimum (0 µg/ml) and maximum (70 µg/ml) observed values. GAM results fit to data from 329 samples with qPCR data. The response variable (*Vc* absolute abundance from qPCR) was log-transformed after adding a pseudocount of 1 and a Gaussian distribution was used. The predictors are ICP1 abundance from qPCR and antibiotics concentration (µg/ml). As with metagenomics data, we fit several GAMs with different combinations of predictors: from a model with all antibiotics and their interaction with ICP1 to separate models with each antibiotic and its interaction with ICP1 and compared them based on their Akaike information criteria (AIC). The most parsimonious model (shown) was retained (the one with the smallest AIC). Both effects of AZI (A) and the ICP1-AZI interaction (B) are significant (Chi-square test, $P<0.05$). Model details are provided in Table S7.

**FIGURE S11**

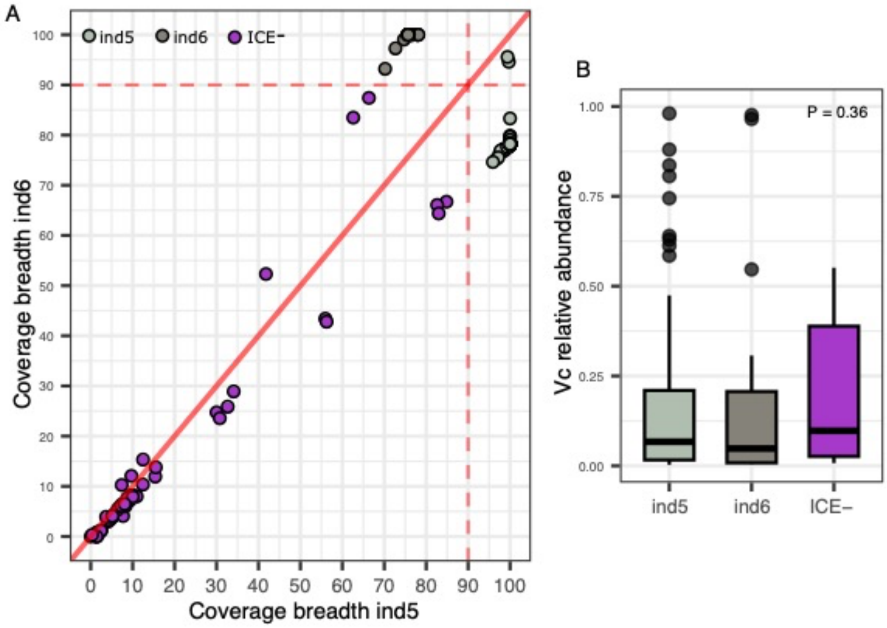

**Fig S11. ICEs are identified unambiguously and are not confounded with *Vc* coverage.** A) Breadth of metagenomic read coverage of the ICEs, ind5 and ind6. Samples were identified as ind5-positive or ind6-positive based on read mapping against reference sequences for *ind5* (NCBI accession GQ463142.1) and *ind6* (accession MK165649.1). The ICE was considered as present when 90% of the reference ICE length was covered by at least one read. Only 2 samples had 100% ind5 and >90% ind6, which were considered ind5-positive. B) Boxplot showing that ICE-negative samples are not associated with lower relative abundance of *Vc*, suggesting that the lack of detecting ICEs is not due to low *Vc* abundance. The *P*-value in panel B is from a Kruskal-Wallis test.

**FIGURE S12**

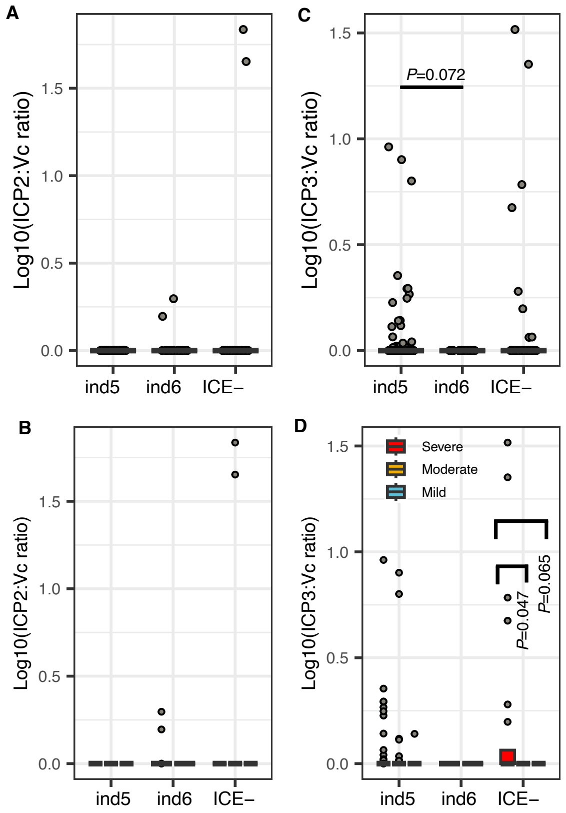

**Fig S12. Ratios of ICP2 and ICP3 to *Vc* across ICE variants**. (A) Distribution of ICP2:*Vc* ratios from metagenomics across patients with different ICE profiles. (B) The same data as (A) binned into boxplots according to dehydration status. (C) Distribution of ICP3:*Vc* ratios across patients with different ICE profiles, (D) The same data as (C) binned into boxplots according to dehydration status. *P*-values are from a Kruskal-Wallis test with Dunn's post-hoc test adjusted with the Benjamini-Hochberg (BH) method. Only *P*-values<0.1 are shown. Only samples with appreciable *Vc* or ICP1 were included (224 samples with *Vc*>0.5% or phages >0.1% of metagenomic reads), of which 54 samples were ICE-, 26 were ind6+ and 144 were ind5+. For clarity, the Y-axes were log10 transformed after adding one to the ratios. Dehydration status is always ordered as severe, moderate, mild, from left to right. The absence of a boxplot means missing data.

**FIGURE S13**

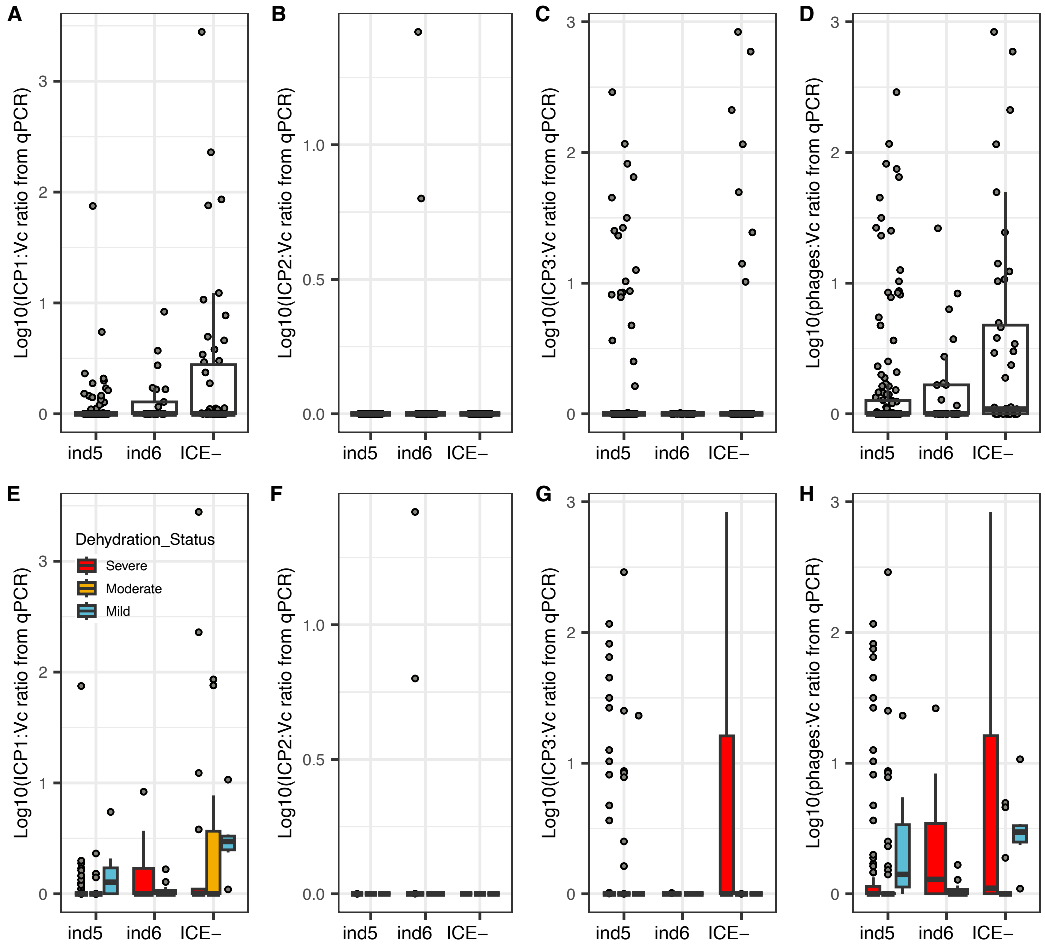

**Fig S13. Integrative conjugative elements (ICEs) are associated with lower ICP1:*V. cholerae* ratios using qPCR data**. (A) Distribution of ICP1:*Vc* ratios across patients with different ICE profiles. (B) The same data as (A) binned into boxplots according to dehydration status. (C) Distribution of ICP2:*Vc* ratios. (D) The same data as (C) binned into boxplots according to dehydration status. (E) Distribution of ICP3:*Vc* ratios, (F) The same data as (E) binned into boxplots according to dehydration status. (G) Distribution of phage:*Vc* ratios, including the sum of all phages (ICP1, ICP2, ICP3). (H) The same data as (G) binned into boxplots according to dehydration status. Only samples with appreciable *Vc* or phage were included (215 samples with *Vc*>0 or phages >0 from qPCR data), of which 50 samples were ICE-, 25 were *ind*6+ and 140 were *ind*5+. The Y-axes were log10 transformed after adding one to the ratios to improve readability. The solid horizontal line is the median and the boxed area is the interquartile range. These results support the findings based on metagenomic data shown in Figures 3 and S12 using qPCR data.

**FIGURE S14**

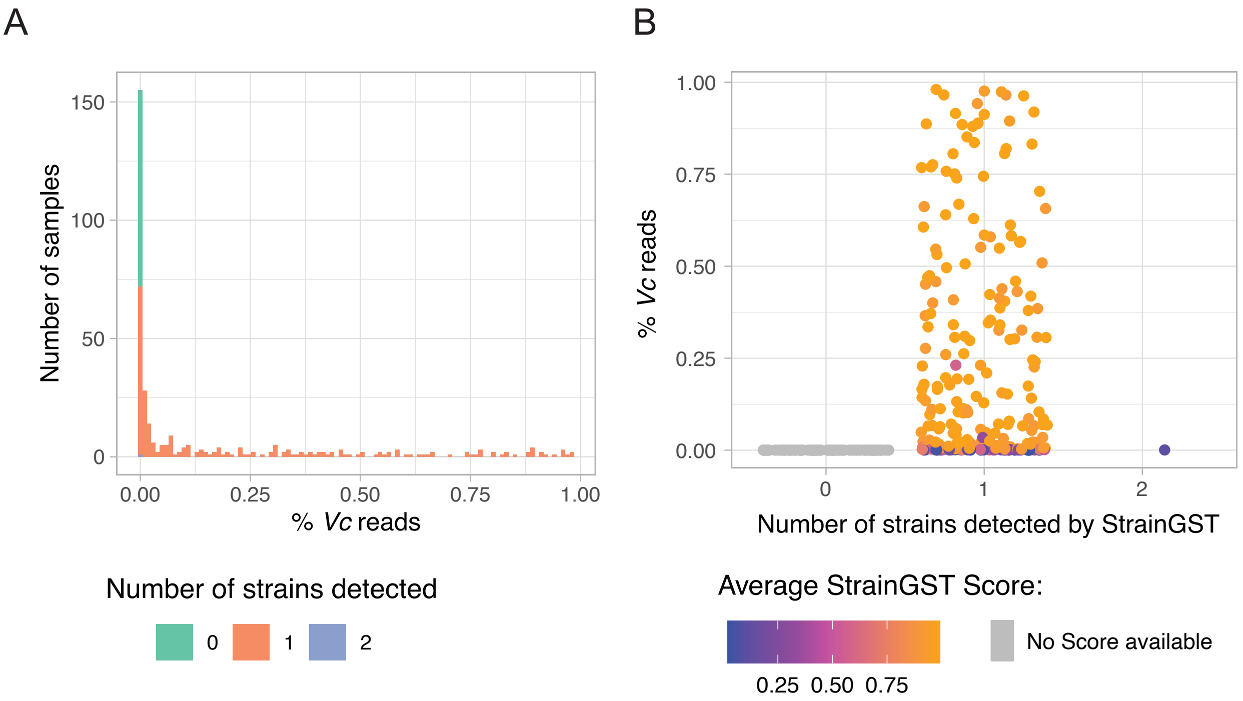

**Fig S14. Mixed infections by more than on *V. cholerae* strain is unlikely in our samples**. (A) The distribution of the number of distinct strains detected across samples. In 260/344 samples, strainGST identified only one reference strain. In 83/344 samples, no reference strain could be identified with confidence, likely due to low coverage of *Vc* (these samples all had <1% *Vc* reads). (B) The relative abundance of *Vc* (% reads) in samples with zero, one, or two strains identified. In one sample, strainGST identified two reference strains, but with a low confidence score and at low *Vc* abundance.

**FIGURE S15**

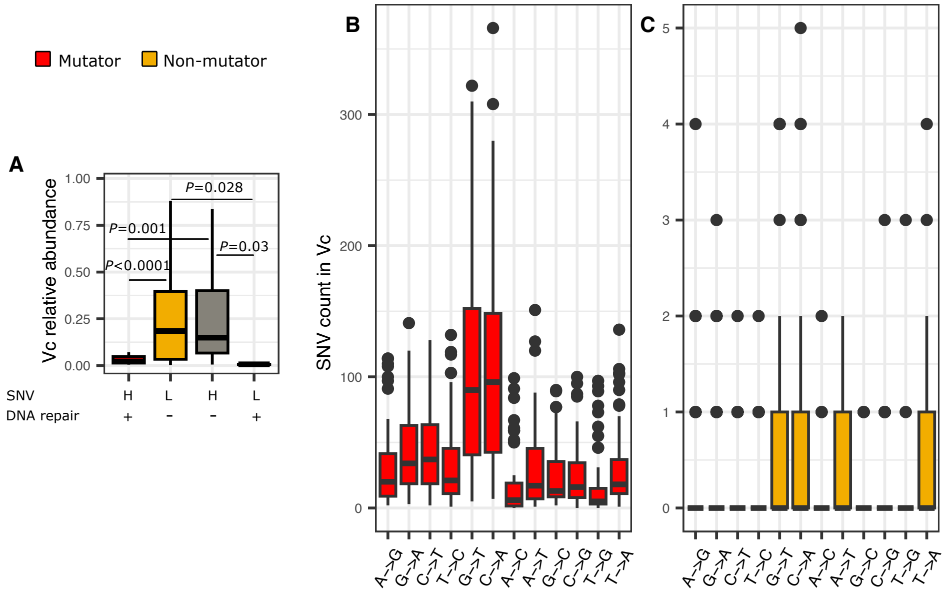

**Fig S15. Characterizing *V. cholerae* hypermutator populations.** A) Low *Vc* relative abundance from metagenomics is associated with DNA repair mutations independently of the number of SNVs. Mutators (red) are defined as having a high (H) number of SNVs (25 or more) in the *Vc* genome, along with one or more nonsynonymous mutations in a DNA repair gene resulting in a predicted defect in DNA repair. Non-mutators (yellow) have neither a high number of SNVs nor a DNA repair defect. *P*-values are from a Kruskal-Wallis test with Dunn's post-hoc test adjusted with the Benjamini-Hochberg (BH) method. Only *P*-values < 0.1 are shown. (B and C) Transversion mutations, particularly G->T and C->A, are more common in mutators compared to non-mutators. Total n=133, with 47 belonging to mutators, 70 to non-mutators, 14 to "high SNVs/no DNA repair defect" and 2 to "low SNVs/with DNA repair defect" groups.

**FIGURE S16**

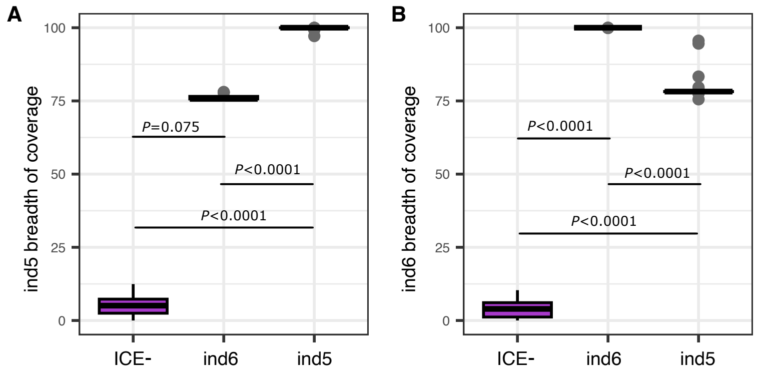

**Fig S16. ICE- are readily distinguishable from *ind5*+ and *ind6*+ samples.** Breadth of ICE coverage across patients with different ICE profiles. (A) Breadth of *ind5* coverage. (B) Breadth of *ind6* coverage. Samples with ambiguous breadth of ICE coverage (in the 40-80% range) were not included in the InStrain analysis and are not included here. *P*-values are from a Kruskal-Wallis test with Dunn's post-hoc test adjusted with the Benjamini-Hochberg (BH) method. n=131 with 24 belonging to ICE-, 18 to *ind6*+, and 89 to *ind5*+ groups.

**FIGURE S17**

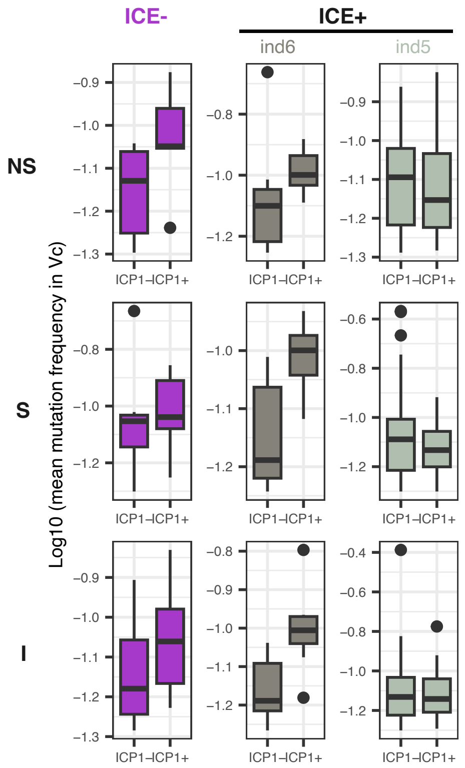

**Fig S17. Boxplots of mutation frequency in *Vc* in the presence or absence of ICP1 and/or ICEs.** Boxplots include 128 samples, of which 41 are ICP1+ (ICP1 detected by qPCR at a Ct cutoff of 28) and 87 are ICP- (Ct below 28). The Y-axes were log10 transformed after adding a pseudocount of 1 to improve readability. The solid horizontal line is the median and the boxed area is the interquartile range. This figure is a version of Fig 4C with qPCR data. NS: nonsynonymous; S: synonymous; I: intergenic SNVs.

**FIGURE S18**

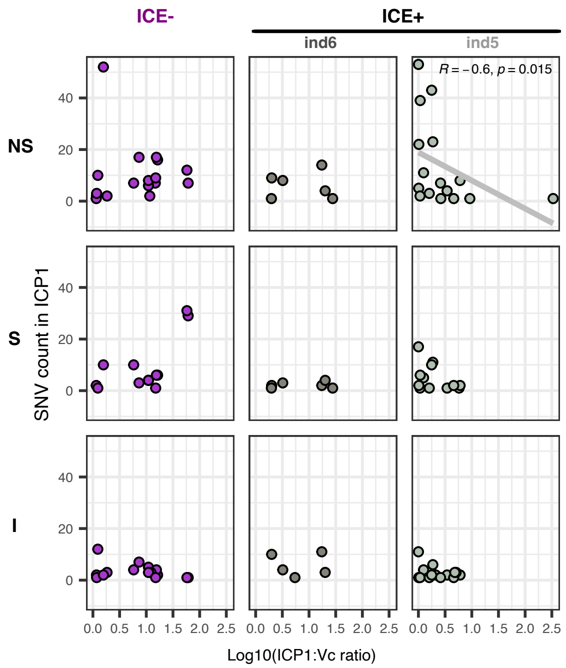

**Fig S18. Number of SNVs in ICP1 as a function of the ICP1:*V.cholerae* ratio from metagenomics and the ICE type encoded by *V. cholerae*.** Single nucleotide variant (SNV) count per sample in the ICP1 genome is plotted as a function of the ICP1:*Vc* ratio for different ICE profiles. All 45 samples for which mutations in ICP1 were called with InStrain were included in this analysis, of which 18 ICE-, 20 ind5 and 7 ind6. Non-synonymous (NS), synonymous (S), and intergenic (I) SNVs are plotted in separate rows. The only significant correlation (Spearman correlation, $P < 0.05$; top right) is shown with the trendline in gray with Spearman correlation coefficient and *P*-value.

**FIGURE S19**

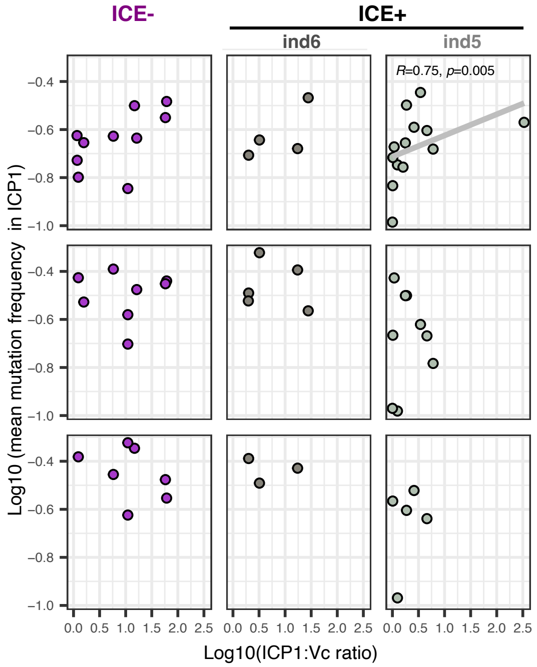

**Fig S19. SNVs in ICP1 as a function of the ICP1:*V.cholerae* ratio from metagenomics and the ICE type encoded by *V. cholerae*.** The mean single nucleotide variant (SNV) frequency per sample in the ICP1 genome is plotted as a function of the ICP1:*Vc* ratio for different ICE profiles. This analysis includes 29 samples for which high frequency mutations in ICP1 were called by InStrain, of which 11 ICE-, 13 *ind5* and 5 *ind6*. Non-synonymous (NS), synonymous (S), and intergenic (I) SNVs are plotted in separate rows. The only significant correlation (Spearman correlation, $P < 0.05$; top right) is shown with the trendline in gray with Spearman correlation coefficient and *P*-value. Only SNV frequencies >0.1 are included.

### SUPPLEMENTARY TABLES

#### TABLE S1

**Table S1. Primers used for PCR and qPCR**

| Target | Primer Name | Sequence 5'-3' | Reference |
|---|---|---|---|
| *ompW (Vibrio cholerae)* | ompW_F<br>ompW_R | CACCAAGAAGGTGACTTTAT<br>GAACTTATAACCACCCGCG | (*76*) |
| *tcpA (Vibrio cholerae)* | *tcpA set2*_F | ACACGATAAGAAAACCGGTCA | (*37*) |
| | *tcpA set2*_R | GCCTTGGTCATATTCTGCGA | |
| ICP1 | gp58.2_F | CAAAGGCAGCAGGTAGGACA | This study |
| | gp58.2_R | CCCTTCAAGCCGTAGTTGGT | |
| ICP2 | gp24_F | AGAAGTCGCAAACGGGGTAC | This study |
| | gp24_R | AACGTGGTTCTCGTGAGTGG | |
| ICP3 | gp19_F | AGACCAACGCCGACTGTTAG | This Study |
| | gp19_R | CGATACCACGGAAAGCCTGT | |
| 16S rDNA1 | Maeda_1048_1067_F | GTGSTGCAYGGYTGTCGTCA | (*60*) |
| | Maeda_1175_1194_R A | ACGTCRTCCMCACCTTCCTC | |

#### TABLE S2

**Table S2. LC MS/MS targets and parameters.** RT= Retention time

| Antibiotic | RT (min) | Extraction Ions |
|---|---|---|
| Azithromycin | 11.4 – 13.1 | 750,591 |
| Ciprofloxacin | 9.5 – 11.2 | 332 |
| Doxycycline | 8.9 – 14.0 | 445,428 |
| Nalidixic Acid | 12.3 – 14.4 | 233 |
| Azithromycin-d5 | 11.4 – 13.1 | 754,595 |
| Ciprofloxacin-d8 | 9.5 – 11.2 | 339 |
| Doxycycline-d5 | 8.9 – 14.0 | 450,433 |
| Metronidazole | 3.5-5.5 | 172 |

**TABLE S3**

**Table S3. Indicator species analysis.** For each group, we report the indicator value (stat) ranging between 0-1, with 1 being the perfect indicator species meaning that the species occurs exclusively in one group. *P*-values are from a permutation test with 9999 iterations as implemented in the indicator species function in R (multipatt from indicspecies library). The analysis was run on all the data (344 samples) and the 37 most dominant species from the PCA in Figure S5. 24 species were selected as indicators of the patient's dehydration status.

| Group | stat | *p* |
|---|---|---|
| **Group 1: mild dehydration** | | |
| *1. Bifidobacterium longum* | 0.372 | 1e-05 |
| *2. Bifidobacterium breve* | 0.304 | 6e-05 |
| *3. Escherichia coli* | 0.266 | 0.00057 |
| *4. Enterococcus aecium* | 0.214 | 0.00437 |
| *5. Bifidobacterium kashiwanohense* | 0.2 | 0.00818 |
| *6. Salmonella enterica* | 0.183 | 0.01262 |
| 7. ICP1 | 0.167 | 0.02641 |
| **Group 2: moderate dehydration** | | |
| *1. Streptococcus pasteurianus* | 0.23 | 0.00272 |
| *2. Roseburia intestinalis* | 0.21 | 0.00596 |
| *3. Streptococcus gallolyticus* | 0.16 | 0.03152 |
| **Group 3: severe dehydration** | | |
| *1. Vibrio cholerae* | 0.333 | 6e-05 |
| *2. Fusobacterium mortiferum* | 0.31 | 0.00011 |
| *3. Faecalibacterium prausnitzii* | 0.291 | 0.00020 |
| *4. Prevotella ruminicola* | 0.284 | 0.00028 |
| *5. Prevotella dentalis* | 0.281 | 0.00030 |
| *6. Brachyspira pilosicoli* | 0.249 | 0.00084 |
| *7. Collinsella aerofaciens* | 0.246 | 0.00149 |
| *8. Bacteroides thetaiotaomicron* | 0.234 | 0.00221 |
| *9. Bacteroides vulgatus* | 0.226 | 0.00309 |
| *10. Prevotella intermedia* | 0.209 | 0.00560 |
| *11. Bacteroides fragilis* | 0.201 | 0.00823 |
| *12. Roseburia [Eubacterium] rectale* | 0.194 | 0.01063 |
| *13.* ICP3 | 0.189 | 0.01050 |
| *14. Roseburia lachnospiraceae bacterium* GAM79 | 0.168 | 0.02409 |

**TABLE S4**

**Table S4. Variables used in the Redundancy analysis (RDA) showing relationships among the seven most dominant bacterial species from PCA (Fig. S5) and explanatory variables**: phages (ICP1, ICP2, ICP3), patient metadata: age in years, vomiting state (yes or no), dehydration status (severe, moderate or mild), the area where the sample was collected, date of sampling, and antibiotic concentration (µg/ml) from quantitative mass spectrometry for Ciprofloxacin (CIP), Doxycycline (DOX) and Azithromycin (AZI). First column: explanatory variables, second column: *P*-values from a permutation test with 999 iterations.

| Variable | *p*[a] |
|---|---|
| CIP | 0.073 |
| DOX | 0.701 |
| AZI | 0.001*** |
| ICP1 | 0.012* |
| ICP2 | 0.336 |
| ICP3 | 0.104 |
| Area_Code | 0.001*** |
| Dehydration status | 0.001*** |
| Collection date | 0.003** |
| Vomiting | 0.002** |
| Age_in_Years | 0.006** |

[a] Permutation test (anova function from the vegan R package).

**TABLE S5**

**Table S5. Antibiotic resistance thresholds.** Minimal inhibitory concentrations (MICs) were established under aerobic or anaerobic conditions in (*16*). Concentrations are in units of µg/ml.

| | **Aerobic** | **Anaerobic** |
|---|---|---|
| Ciprofloxacin | 0.016 | 0.063 |
| Azithromycin | 1 | 8 |
| Doxycycline | 0.13 | 0.13 |

### TABLE S6

**Table S6. Generalized additive models included in the model selection using metagenomic data**. GAMs were fit with *Vc* relative abundance as a function of ICP1 relative abundance, antibiotic concentration (µg/ml) and their interactions. The selection was based on ∆AIC. We report ∆AIC, R syntax for the formula, the predictors (fixed effects) and the corresponding *P*-values (Chi-square test), as well as the *P*-value corresponding to dehydration random effect (RE) and the adjusted r-squared of the model. nt: not tested.

| model | ∆AIC | Formula | ICP1 | AZI | ICP1*AZI | Dehyd (RE) | R2 |
|---|---|---|---|---|---|---|---|
| Mod1 (selected) | 0 | Vc ~ s(ICP1) + s(AZI) + te(ICP1, AZI) + s(Dehydration_Status, bs = "re") | 0.614 | 0.002 | 0.026 | 3.66e-07 | 0.031 |
| Mod2 | 1.3 | Vc ~ s(ICP1) + s(AZI) + s(Dehydration_Status, bs = "re") | 0.077 | 0.004 | Not tested | 2.76e-07 | 0.035 |
| Mod3 | 3.9 | Vc ~ s(ICP1) + s(AZI) + s(CIP) + s(DOX) + te(ICP1, AZI) + te(ICP1, CIP) + te(ICP1, DOX) +s(Dehydration_Status, bs = "re") | 0.606 | 0.002 | 0.025 | 4.08e-07 | 0.027 |
| Mod4 | 5.2 | Vc ~ s(ICP1) + s(AZI) + s(CIP) + s(DOX) + s(Dehydration_Status, bs = "re") | 0.078 | 0.003 | nt | 3.06e-07 | 0.032 |
| Mod5 | 11.7 | Vc ~ s(ICP1) + s(DOX) + te(ICP1, DOX) + s(Dehydration_Status, bs = "re") | 0.059 | nt | nt | 3.65e-08 | 0.035 |
| Mod6 | 11.9 | Vc ~ s(ICP1) + s(CIP) + te(ICP1, CIP) + s(Dehydration_Status, bs = "re") | 0.664 | nt | 4.35e-08 | 4.35e-08 | 0.022 |
| Mod7 | 12.4 | Vc ~ s(ICP1) + s(AZI) + te(ICP1, AZI, by = Dehydration_Status) | | | | | |
| Mod1(B)[a] | 29.3 | Vc ~ s(ICP1) + s(AZI) + te(ICP1, AZI) | 0.914 | 0.003 | 0.015 | nt | -0.017 |

[a] Without dehydration RE

**TABLE S7**

**Table S7. Generalized additive models included in the model selection using qPCR data.** Generalized additive models included in the model selection. GAMs were fit with *Vc* as a function of ICP1 (absolute abundances from qPCR with Ct threshold=28), antibiotic concentration (µg/ml) and their interactions, the same methods as for the models fit with metagenomics data were used. The selection was based on ∆AIC. We report ∆AIC, R syntax for the formula, the predictors (fixed effects) and the corresponding P-values (Chi-square test), as well as the *P*-value corresponding to dehydration random effect (RE) and the adjusted r-squared of the model. nt: not tested.

| **Model** | **∆AIC** | **Formula** | **ICP1** | **AZI** | **ICP1*AZI** | **Dehyd(RE)** | **R2** |
|---|---|---|---|---|---|---|---|
| Mod1 | 0 | log_qpcr_vc ~ s(qPCR_ICP1_PFUml_CT28) + s(AZI) + s(CIP) + s(DOX) + te(qPCR_ICP1_PFUml_CT28, AZI) +te(qPCR_ICP1_PFUml_CT28, CIP) + te(qPCR_ICP1_PFUml_CT28, DOX) + s(Dehydration_Status,bs = "re") | 0.622 | 0.001 | 0.017 | < 2e-16 | 0.172 |
| Mod2 | 0.4 | log_qpcr_vc ~ s(qPCR_ICP1_PFUml_CT28) + s(AZI) + s(CIP) + s(DOX) + s(Dehydration_Status, bs = "re") | 0.023 | 0.001 | nt | < 2e-16 | 0.172 |
| Mod3 | 0.9 | log_qpcr_vc ~ s(qPCR_ICP1_PFUml_CT28) + s(AZI) + te(qPCR_ICP1_PFUml_CT28, AZI) + s(Dehydration_Status, bs = "re") | 0.636 | 0.001 | 0.015 | < 2e-16 | 0.163 |
| Mod4 | 1.3 | log_qpcr_vc ~ s(qPCR_ICP1_PFUml_CT28) + s(AZI) + s(Dehydration_Status, bs = "re") | 0.018 | 0.001 | nt | < 2e-16 | 0.163 |
| Mod5 | 5.3 | log_qpcr_vc ~ s(qPCR_ICP1_PFUml_CT28) + s(AZI) + te(qPCR_ICP1_PFUml_CT28, AZI, by = Dehydration_Status) | 0.265 | 0.004 | <0.003 * | nt | 0.169 |
| Mod6 | 11.8 | log_qpcr_vc ~ s(qPCR_ICP1_PFUml_CT28) + s(DOX) + te(qPCR_ICP1_PFUml_CT28, DOX) + s(Dehydration_Status, bs = "re") | 0.019 | nt | nt | < 2e-16 | 0.136 |
| Mod7 | 11.9 | log_qpcr_vc ~ s(qPCR_ICP1_PFUml_CT28) + s(CIP) + te(qPCR_ICP1_PFUml_CT28, CIP) + s(Dehydration_Status, bs = "re") | 0.786 | nt | nt | < 2e-16 | 0.134 |
| Mod8 (Mod3 without RE) | 30.6 | log_qpcr_vc ~ s(qPCR_ICP1_PFUml_CT28) + s(AZI) + te(qPCR_ICP1_PFUml_CT28, AZI) | 0.393 | 0.0001 | 0.031 | nt | 0.079 |

*For all dehydration status levels

**TABLE S8**

**Table S8. Generalized linear mixed models (**GLMMs) **included in the model selection.** The response is SNV count in *Vc* and the predictors are ICP1 relative abundance from metagenomics, antibiotic concentration (µg/ml) and their interaction. We report ∆AIC, the R syntax for the formula, the predictors, and the corresponding *P*-values (Wald test). As an evaluation of the goodness of fit, we report the adjusted R-squared (R2) and *P*-value from the comparison of the models and null models (exactly the same model but with no fixed terms) (likelihood-ratio test, anova function from stats package in R). Antibiotics terms are not shown (not significant, *P*>0.05, Wald test). nt: not tested.

| **Model** | **dAIC** | **Formula** | ***Vc*** | **ICP1** | **ICP1*Vc** | **R2** | **P/null model** |
|---|---|---|---|---|---|---|---|
| Mod1 (Selected) | 0.0 | SNV_nb ~ Vc + Vc:ICP1 | 0.002 | nt | 0.004 | 0.377 | 7e-04 |
| Mod2 | 0.6 | SNV_nb ~ ICP1 + Vc + Vc:ICP1 | 0.007 | 0.266 | 0.012 | 0.408 | 0.001 |
| Mod3 | 1.6 | SNV_nb ~ ICP1 + Vc + AZI + Vc:ICP1 | 0.008 | 0.267 | 0.015 | 0.409 | 0.002 |
| Mod4 | 3.2 | SNV_nb ~ ICP1 + Vc + CIP + AZI + Vc:ICP1 | 0.007 | 0.273 | 0.015 | 0.412 | 0.004 |
| Mod5 | 5.0 | SNV_nb ~ Vc | 0.002 | nt | nt | 0.228 | 0.006 |
| Mod6 | 5.2 | SNV_nb ~ ICP1 + Vc + CIP + DOX + AZI + Vc:ICP1 | 0.00 | 0.273 | 0.015 | 0.412 | 0.008 |
| Mod6 | 5.9 | SNV_nb ~ ICP1 + Vc + CIP + DOX + AZI + ICP1:CIP + ICP1:AZI + Vc:ICP1 | 0.0428 | 0.163 | 0.038 | 0.458 | 0.008 |
| Mod7 | 7.0 | SNV_nb ~ ICP1 + Vc + CIP + AZI + DOX + Vc:ICP1:CIP + Vc:ICP1:AZI + Vc:ICP1:DOX | 0.015 | 0.732 | nt | 0.442 | 0.012 |
| Mod8 | 7.1 | SNV_nb ~ ICP1 + Vc + CIP + DOX + AZI + ICP1:AZI + Vc:ICP1 | 0.047 | 0.275 | 0.059 | 0.422 | 0.015 |
| Mod9 | 12.1 | SNV_nb ~ ICP1 | nt | 0.506 | nt | 0.018 | 0.527 |

**TABLE S9**

**Table S9. Generalized additive models (GAMs) included in the model selection.** The response is the average frequency of NS mutations in *Vc* and the predictors are ICP1 (relative abundance from metagenomics), antibiotic concentration in µg/ml (antbx), ICE presence/absence and mutation type (mut) (NS: non-synonymous, S:synonymous and I:intergenic) as well as their interactions. We defined a variable (ICE.by.mut) as the combination of ICE and mutation type with 9 levels (mutation type*ICE) to represent the interaction between mutation types (NS, S, I) and ICE variants (*ind5*, *ind6*, ICE-). Model selection was based on ∆AIC. We report ∆AIC, the R syntax for the formula, the predictors, and the corresponding *P*-values (Chi-square test) and the adjusted R-squared of the model. Our data was insufficient to fit models with the interaction between ICP1 and antibiotics. nt: not tested.

| | ∆AIC | **formula** | ***P* (ICP1*ICE*mut)** | | | ***P* (antbx*ICE* mut)** | **R2** |
|---|---|---|---|---|---|---|---|
| | | | Ind5*NS | Ind6*NS | ICE-*NS | | |
| Mod1 | 0.0 | mean_freq ~ s(ICP1, by = ICE.by.mut) | 0.1336 | 0.642 | 0.0469 | nt | 0.021 |
| Mod2 | 10.3 | mean_freq ~ s(ICP1, by = ICE.by.mut) + s(AZI, by = ICE.by.mut) | 0.1338 | 0.9408 | 0.0373 | >0.05 | 0.01 |
| Mod3 | 23.4 | mean_freq ~ s(ICP1, by = ICE.by.mut) + s(AZI, by = ICE.by.mut) + s(CIP, by = ICE.by.mut) | 0.1142 | 0.9199 | 0.0411 | >0.05 | -0.011 |
| Mod4 | 30.6 | mean_freq ~ s(ICP1, by = ICE.by.mut) + s(AZI, by = ICE.by.mut) + s(CIP, by = ICE.by.mut) + DOX * ICE.by.mut | 0.11048 | 0.87574 | 0.04109 | >0.05 | 0.0313 |
| Mod5 | | mean_freq ~ s(ICP1, by = ICE.by.mut) + s(AZI, by = ICE.by.mut) + s(CIP, by = ICE.by.mut) + DOX * ICE.by.mut + te(ICP1,AZI, by = ICE.by.mut) + te(ICP1, CIP, by = ICE.by.mut) + DOX:ICP1:ICE.by.mut | Not enough data to fit | | | | |

### TABLE S10

**Table S10. Most frequently mutated *Vc* genes in samples with high ICP1:*V.cholerae* ratios.** Mutation count denotes the total number of SNVs and patient count denotes the number of patients in which a mutation was observed. Genes were identified with prodigal v2.6.3 and annotated with eggNOG-Mapper v2 as well as manually with blastx against NCBI database within the Basic Local Alignment Search Tool (BLAST) online tool. Genes that contain mutations in samples with ICP1>*Vc* exclusively are shown in bold. Only the top 15 genes are shown, See Data File S3 for complete list.

| COG (eggnog) | PFAM (eggnog) annotation | NCBI annotation | Mutation count | Patient count |
|---|---|---|---|---|
| Q | HemolysinCabind_Peptidase_M10_C | retention module-containing protein WP_001191814.1 | 5 | 2 |
| **M** | **Peptidase_S13** | **D-alanyl-D-alanine carboxypeptidase/D-alanyl-D-alanine-endopeptidase (genbank : AAF93798.1)** | **4** | **1** |
| V | ACR_tran | multidrug resistance protein, putative AAF93795.1 | 4 | 1 |
| O | PPC_Peptidase_M9_Peptidase_M9_N | TPA: collagenase HAS4622795.1 | 4 | 1 |
| P | BPD_transp_1 | ABC transporter permease subunit WP_000252168.1 | 4 | 1 |
| **K** | **MerR_1** | **MerR family transcriptional regulator WP_000226962.1** | **3** | **1** |
| C | CCG_FADoxidase_C_FAD_binding_4_Fer4_7_Fer4_8 | FAD-binding and (Fe-S)-binding domain-containing protein WP_000188699.1 | 3 | 1 |
| **C** | **Gp_dh_C_Gp_dh_N** | **glyceraldehyde 3-phosphate dehydrogenase GenBank: AAF96741.1** | **3** | **1** |
| S | DUF3302 | DUF3302 domain-containing protein WP_000478180.1 | 2 | 1 |
| IQ | PPbinding | Chain A, 3-oxoacyl-[acyl-carrier-protein] synthase 2 PDB: 4JRH_A | 2 | 1 |
| **C** | **Fer4_12_Radical_SAM** | **methyl-accepting chemotaxis protein WP_000383592.1** | **2** | **2** |
| Q | FtsX_MacB_PCD | ABC transporter permease WP_000645916.1 | 2 | 1 |
| **C** | **CCG_Fer4_8** | **anaerobic glycerol-3-phosphate dehydrogenase subunit GlpC WP_001014995.1** | **2** | **1** |
| GM | CoA_binding_3_Polysacc_synt_2 | nucleoside-diphosphate sugar epimerase/dehydratase WP_000494952.1 | 2 | 1 |
| F | PpxGppA | guanosine-5'-triphosphate,3'-diphosphate diphosphatase WP_000076046.1 | 2 | 1 |

### TABLE S11

**Table S11. Most frequently mutated *Vc* genes in samples with low ICP1:*V.cholerae* ratios.** Mutation count denotes the total number of SNVs and patient count denotes the number of patients in which a mutation was observed. Genes were identified with prodigal v2.6.3 and annotated with eggNOG-Mapper v2 as well as manually with blastx against NCBI database within the Basic Local Alignment Search Tool (BLAST) online tool. Genes that contain mutations in samples with ICP1<*Vc* exclusively are shown in bold. Only the top 15 genes are shown, see Data File S4 for complete list.

| COG category (eggnog) | PFAM (eggnog) annotation | NCBI annotation | Mutation count | Patient count |
|---|---|---|---|---|
| O | PPC_Peptidase_M9_Peptidase_M9_N | TPA: collagenase HAS4622795.1 | 56 | 15 |
| **MQ** | **ACD_ADPrib_exo_Tox_Anthrax_toxA_Hydrolase_4_MLD_Peptidase_C80_RtxA** | **TPA: MARTX multifunctional-autoprocessing repeats-in-toxin holotoxin RtxA HAS4620517.1** | **14** | **6** |
| **PT** | **GGDEF_Hemerythrin** | **GGDEF domain-containing protein WP_001190450.1** | **12** | **4** |
| **T** | **EAL_GGDEF** | **EAL domain-containing protein WP_000160066.1** | **11** | **2** |
| **S** | **Glyco_hydro_129** | **hypothetical protein VC_A0254 AAF96165.1** | **11** | **2** |
| **S** | **Betaprism_lec_Hemolysin_N_Leukocidin** | **Cytolysin vcc (ncbi) WP_001125271.1 (hlyA gene) [Levade et al 2021]** | **10** | **3** |
| C | FixO | cytochrome-c oxidase, cbb3-type subunit II WP_000097777.1 | 9 | 1 |
| **L** | **Fapy_DNA_glyco_H2TH_zfFPG_IleRS** | **bifunctional DNA-formamidopyrimidine glycosylase/DNA-(apurinic or apyrimidinic site) lyase WP_001114647.1** | **9** | **3** |
| I | AcylCoA_dh_1_AcylCoA_dh_M_AcylCoA_dh_N_DUF1974 | acyl-CoA dehydrogenase FadE WP_000404358.1 | 8 | 4 |
| L | Phage_int_SAM_4 | hypothetical protein WP_000222725.1 | 8 | 4 |
| M | ACD_PG_binding_1_Pesticin_Phage_GPD | Type VI secretion system tip protein VgrG WP_000212125.1 | 7 | 1 |
| K | LacI_Peripla_BP_3 | DNA-binding transcriptional regulator CytR WP_000224452.1 | 7 | 1 |
| T | EAL | EAL domain-containing protein WP_000547542.1 | 7 | 2 |
| E | SDH_alpha_SDH_beta | sodium:solute symporter WP_001882525.1 | 7 | 4 |
| S | Inovirus_Gp2 | inovirus Gp2 family protein WP_000149629.1 | 7 | 4 |

### TABLE S12

**Table S12. Most frequently mutated ICP1 genes in samples with ICE *ind5* detected.** Mutation count denotes the total number SNVs (NS SNVs with frequency>0.1) and patient count denotes the number of patients in which a mutation was observed. Only samples with the ICE *ind5* are included (13 samples out of the 27 samples for which high frequency NS mutations were called in ICP1 with InStrain). Genes were annotated manually with blastx against NCBI database within the Basic Local Alignment Search Tool (BLAST) online tool. Most of the genes had no hit with eggNOG-Mapper. Only the top 15 genes are shown; see Data File S5 for the complete list,

| NCBI annotation | Mutation count | Patient count |
|---|---|---|
| hypothetical protein ViPhICP1_gp184 YP_004251125.1 | 33 | 3 |
| hypothetical protein ViPhICP1_gp147 YP_004251088.1 | 5 | 2 |
| anaerobic ribonucleotide reductase small subunit YP_004251055.1 | 4 | 3 |
| hypothetical protein ViPhICP1_gp150 YP_004251091.1 | 4 | 2 |
| acyl carrier protein YP_004251038.1 | 4 | 3 |
| hypothetical protein ViPhICP1_gp226 YP_004251167.1 | 3 | 3 |
| nicotinamide phosphoribosyl transferase YP_004251171.1 | 3 | 3 |
| hypothetical protein ViPhICP1_gp139 YP_004251080.1 | 2 | 1 |
| hypothetical protein ViPhICP1_gp166 YP_004251107.1 | 2 | 2 |
| hypothetical protein ViPhICP1_gp018 YP_004250959.1 | 2 | 2 |
| hypothetical protein ViPhICP1_gp175 YP_004251116.1 | 2 | 1 |
| hypothetical protein ViPhICP1_gp199 YP_004251140.1 | 2 | 2 |
| ribonucleoside diphosphate reductase, beta chain YP_004251147.1 | 2 | 2 |
| hypothetical protein ViPhICP1_gp068 YP_004251009.1 | 2 | 2 |
| hypothetical protein ViPhICP1_gp070 YP_004251011.1 | 2 | 1 |
